# The evolution of stage-specific virulence: differential selection of parasites in juveniles

**DOI:** 10.1101/324632

**Authors:** Ryosuke Iritani, Elisa Visher, Mike Boots

**Affiliations:** Biosciences, College of Life and Environmental Science, University of Exeter, Exeter, United Kingdom.; Department of Integrative Biology, University of California, Berkeley, 3040 Valley Life Sciences Building #3140, Berkeley, CA 94720-3140.

**Keywords:** Adaptive dynamics, Age-structured population, Life-history evolution, Parasite virulence, Senescence

## Abstract

The impact of infectious disease is often very different in juveniles and adults, but theory has focused on the drivers of stage-dependent defense in hosts rather than the potential for stage-dependent virulence evolution. Stage-structure has the potential to be important to the evolution of pathogens because it exposes parasites to heterogeneous environments in terms of both host characteristics and transmission routes. We develop a stage-structured (juvenile-adult) epidemiological model and examine the evolutionary outcomes of stage-specific virulence under the classic assumption of a transmission-virulence trade-off. We show that selection on virulence against adults remains consistent with the classic theory. However, the evolution of juvenile virulence is sensitive to both demography and transmission pathway with higher virulence against juveniles being favored either when the transmission pathway is assortative (juveniles preferentially interact together) and the juvenile stage is short, or in contrast when the transmission pathway is disassortative and the juvenile stage is long. These results highlight the potentially profound effects of host stage-structure on determining parasite virulence in nature. This new perspective may have broad implications for both understanding and managing disease severity.

**Impact summary:** Understanding the evolution of parasite virulence remains one of the most important questions in evolutionary ecology. Virulence is often very different in young and old hosts, but previous theory has presumed that these differences are attributed to adaptation in host defense rather than parasite adaptation. However, stage-structure within host populations can expose parasites to heterogeneous environments, which may lead to differential selection on parasite virulence (stage-specific virulence). Surprisingly, no study has investigated the effects of hosts’ stage-structure on the evolution of stage-specific virulence. We present a theoretical analysis to examine when selection can favor higher virulence against juveniles (juvenile-virulence) versus adults (adult-virulence). Our key result is that higher juvenile-virulence is selected for either when the transmission is assortative within age classes and maturation is slow, or when the transmission is disassortative (occurring predominantly between-classes) and maturation is relatively fast. These at first sight contrasting outcomes can be understood as adaptation to the exploitation of the more available host stage. Although the data on assortativity in infectious disease systems is limited, empirical studies for the virulence of Great Island Virus in guillemots (*Uria aalge*) and for salmon louse in pink salmon (*Oncorhynchus gorbuscha*) are consistent with our predictions. Our work provides testable predictions for stage-specific virulence and presents a novel mechanism that may explain variation in virulence in nature. There are also management implications for conservation, public health, vaccination programs, and farming to understanding the drivers of stage dependent virulence.

## Introduction

Understanding how parasites are selected to exploit their hosts remains a central research question in the evolutionary ecology of host-parasite interactions (Smith 1904; Ball 1943; Anderson & May 1982; Read 1994; Ebert & Herre 1996; Frank 1996; Alizon *et al.* 2009; Schmid-Hempel 2011; Bull & Lauring 2014; Cressler *et al.* 2016), with important implications for host persistence (Boots & Sasaki 2003; De Castro & Bolker 2005), disease management (Dieckmann 2005), and host-parasite coevolution (Boots *et al.* 2009). Although most of the theory of evolution of virulence (defined in this literature as the increased death rate due to infection) focuses on homogeneous host populations, heterogeneity within host populations is ubiquitous in nature (Anderson & May 1992, Chapter 8-11). One typical form of host heterogeneity is stage-related structure (e.g., juveniles and adults), and a number of recent ecological studies have examined the impacts of host populations’ stage-related heterogeneity on disease epidemiology (e.g., Dwyer 1991; Fleming-Davies *et al.* 2015; Hite *et al.* 2016; for theory, Ashby & Bruns 2018). In these studies, the differences in virulence across life stages have been explained as age-related variation in tolerance, resistance, exposure, immunocompetence, and susceptibility and affected by maternal and acquired immunity (Hudson & Dobson 1995; Wilson *et al.* 2002).

In addition to age-related variation in the hosts, different host age-classes expose parasites to specific environmental heterogeneity (see Ashby & Bruns 2018, for theory). Given this, parasites may adaptively tune conditional exploitation against certain stage classes, e.g., through plasticity or by infecting tissues and/or cells that are differentially expressed at different stage classes. In principle, stage-specific virulence may occur as a result of parasite adaptation in two ways. First, stage-structure can generate different infectious periods, for instance due to the substantial difference in natural mortality between juveniles and adults (Jones *et al.* 2013). Infectious period is therefore stage-dependent, which, according to the theory (Day 2001; Gandon *et al.* 2001; Day & Proulx 2004; Gandon 2004; Alizon *et al.* 2009; Cressler *et al.* 2016), may induce selection on virulence such that a shorter infectious period in a certain stage of hosts favors higher virulence. Secondly, the hosts’ stage-structure can generate biased transmission pathways, thereby exposing the parasites to temporal heterogeneity, which may induce additional selective pressures (reviewed in Lion & Metz 2018). For instance, spatiotemporal segregation between juveniles and adults, which is typical of humans (Rohani *et al.* 2010), amphibians (Kilpatrick *et al.* 2010), and insects (Briggs & Godfray 1995), can produce assortative transmission pathway (i.e., juvenile-juvenile and adult-adult transmission might be more likely than juvenile-adult transmission), such that parasites infecting a certain stage of hosts are likely to be transmitted to the same stage of hosts. The assortativity in transmission can thus facilitate or limit the access of parasites to differential quality of resources of hosts. Despite these potentially important selective forces on stage-specific virulence, the implications of host stage-structure for parasite fitness and evolution of stage-specific virulence have not been examined.

Here, we extend classic models of virulence evolution to include two host stage-classes (pre and post reproductive) where the juveniles mature into adults, the adults reproduce, and transmission between the stage-classes is characterized by a matrix of transmission pathways. We explore the evolutionary outcomes of stage-specific virulence in light of classic theory of life history evolution in heterogeneous populations. We use the adaptive dynamics toolbox (Hofbauer & Sigmund 1990; Dieckmann & Law 1996), assessing the joint evolution of virulence against adults (adult-juvenile) and that against juveniles (juvenile-virulence). The models show that selection for virulence can differ in juveniles compared to adults.

## Method

We consider a host population structured into juvenile (J) and adult (A) stages, in which juveniles are by definition incapable of reproduction. The density of susceptible or infected juveniles is denoted *S*_J_ or *I*_J_ respectively, and that of susceptible or infected adults is denoted *S*_A_ or *I*_A_ respectively. Combining an epidemiological SI-model with a stage-structured model yields the following ordinary differential equations (ODEs; Appendix A1):

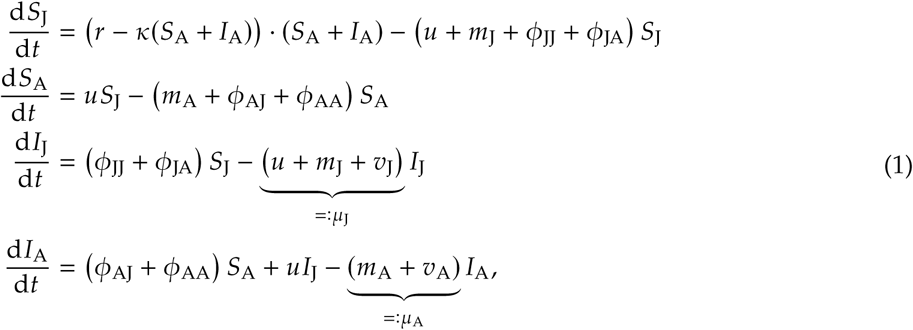

where: *r* represents a fecundity of the adult hosts per capita (assumed to be the same for susceptible and infected adults), which is reduced by a density-dependent factor *κ*; juveniles mature into adults at a rate *u*; *m*_X_ represents the background mortality for a stage-X host; *ϕ*_XY_ represents the rate at which a susceptible stage-X host gets transmitted from an infected stage-Y host (i.e., force of infection from infectious Y to susceptible X per capita, Fig 2; also see below for formula); *v*_J_ (or *v*_A_) represents the virulence against a juvenile (or adult) host. The reciprocal of the infectious period for juveniles (or adults) is given by *µ*_J_ = *u* + *m*_J_ + *v*_J_ (or by *µ*_A_ = *m*_A_ + *v*_A_ respectively). For alternative approaches including physiologically structured population modeling and infection-age modeling with continuous stage structure, see Day *et al.* (2011), Mideo *et al.* (2011), and de Roos & Persson (2013).

Maturation and natural mortality can both affect the relative length of the adult stage. To quantify this, let *θ*_A_ be the expected fraction of time a host individual spends as an adult in the entire lifespan in the absence of disease, which reads (Appendix A2):

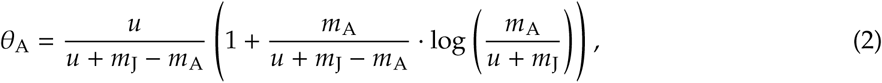

from which we can check that extremely slow (or fast) maturation, *u* → 0 (or *u* → +∞ respectively), leads to *θ*_A_ → 0 (or 1 respectively; note, a special case for *u* + *m*_J_ = *m*_A_ yields *θ*_A_ = *u*/ (2(*u* + *m*_J_)); Appendix A2). We use *θ*_A_ as a characteristic parameter of the stage-structured host populations.

The force of infection for a stage-X host from a stage-Y host (with X and Y running across J and A) in Eqn (1) involves with three processes: susceptibility *α*_X_ (the likeliness for which a stage-X host becomes infected, given a reception of pathogen propagule), transmission pathway *σ*_XY_ (which represents the probability that a pathogen propagule, given that it was produced within Y-stage host, is transferred to a X-stage host), and infectiousness *β*_Y_ (the propagule-production from a stage-Y host; see Fig 2 A):

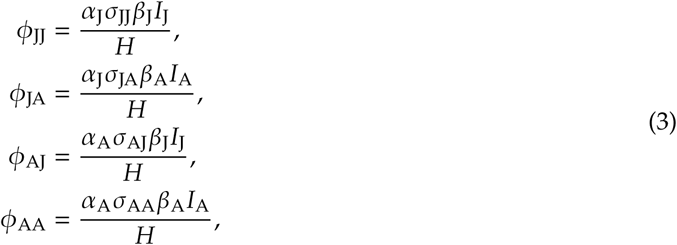

with *H* = *S*_J_ + *I*_J_ + *S*_A_ + *I*_A_ the total density of hosts, such that the transmission is frequency-dependent, as is assumed in previous studies of stage-structured epidemiological dynamics (e.g., Bernhauerová 2016). Also, to link virulence and transmission, we use the trade-off relationship given by 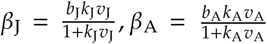, where *k*_X_ tunes the efficiency of virulence for infectiousness from stage-X hosts, and *b*_X_ represents the upper bound of infectiousness from stage-X hosts. Note that we assumed that transmission is a concave function of virulence to restrict our primary attention to stable evolutionary outcomes (Otto & Day 2007).

For quantifying the structure of transmission pathways with a single parameter, we first assume symmetricity, *σ*_AJ_ = *σ*_JA_ (“consistency condition” for mixing structure; Diekmann *et al.* 2012, Chapter 12). Also, we normalize the system, and assume that *σ*_AA_ = 1 – *σ*_AJ_ = 1 – *σ*_JA_ = *σ*_JJ_. With these assumptions, the assortativity measure is given by:

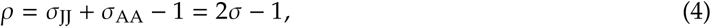

where *ρ* varies from –1 to 1 (and note that these assumptions on the symmetric, normalized transmission pathways will be relaxed in subsequent analyses). If –1 ≤ *ρ* < 0, then within-stage transmission is less likely compared to between-stage transmission (such a structure of transmission pathways is said to be “disassortative”). Instead, if 0 < *ρ* ≤ 1, then within-stage transmission is more likely than between-stage transmission (assortative transmission). *ρ* = 0 indicates that transmission is unbiased (“random” transmission). In the extreme case, *ρ* = 1 (or –1) indicates that transmission occurs exclusively within the stages (or between the stages, respectively; Fig 2B-D). For a more general treatment of contact structure, see Brauer & Castillo-Chavez (2012) (Chapter 3-5) and Diekmann *et al.* (2012) (Chapter 12).

We use the adaptive dynamics toolbox (Hofbauer & Sigmund 1990; Dieckmann & Law 1996) to study the long-term evolutionary dynamics of stage-specific virulence. Throughout the paper we assume that parasites show stage-specific virulence, *v*_J_ and *v*_A_, with no association or correlation between them (i.e., we study the joint evolutionary dynamics of (*v*_J_, *v*_A_)). First, suppose that the system of ODEs in Eqn (1) has reached an endemic equilibrium: 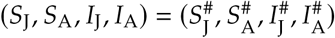 for a given (or wild type) virulence **v** := (*v*_J_, *v*_A_) (where the symbol := will be henceforth used for defining a quantity). We then introduce a rare mutant 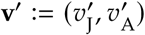 attempting to invade a monomorphic wild type virulence **v**, assuming weak selection (|**v**′ – **v**| is very small). For more details, see Appendix A3.

To assess the possibility of mutant invasion, we define the invasion fitness, denoted *w*, by using the Next-Generation Theorem (van den Driessche & Watmough 2002; Hurford *et al.* 2010). The “next-generation matrix” (that determines the long term growth of the mutant, denoted **G**′) can be written as the product of five matrices:

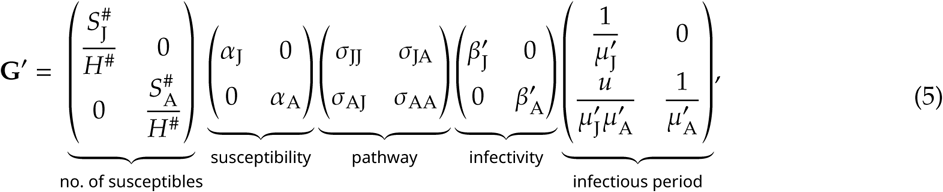

where 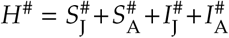 (the total density of the hosts at the endemic equilibrium), 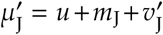 (the reciprocal of the infectious period of juveniles infected by the mutant), and 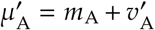 (the reciprocal of the infectious period of adults infected by the mutant). Eqn (5) offers a natural interpretation of the reproductive success of the mutant by partitioning the epidemiological process, in agreement with models of transmission dynamics in heterogeneous host populations (Craft 2015; VanderWaal & Ezenwa 2016; White *et al.* 2017). The first matrix represents the availability of susceptible hosts, each with a specific susceptibility (the second matrix); the third matrix represents the transmission pathways across stages; the fourth matrix represents the infectiousness of infected hosts per capita, and the fifth matrix represents the stage-specific infectious period; the left bottom element represents a conditional, expected infectious period of adult hosts, given by the maturation probability of a juvenile infected by the mutant 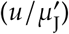 times the expected infectious period of adult hosts 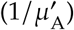.

The invasion fitness is determined by the dominant eigenvalue of **G**′ (denoted Λ[**G**′]), which turns out to exhibit a complicated expression. We therefore use a simpler (but equivalent) measure for the invasion fitness:

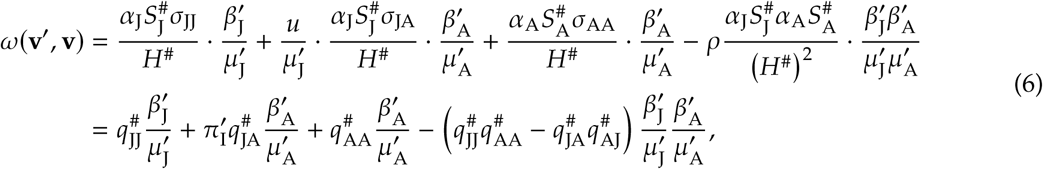

with shorthand notation 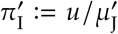 (the probability of maturation of juveniles infected by the mutant) and 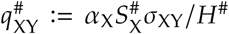 (the availability of X-stage hosts to the parasites infecting a Y-stage host per propagule-production). The factor 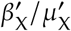 represents the total number of propagule produced by a X-stage host infected by the mutant. We find that the condition for which the mutant outcompetes the wild type *ω*(**v**′, **v**) > 1 holds if and only if Λ[**G**′] > 1, under weak selection (see Appendix A4).

The initial phases of evolutionary dynamics are determined by the selection gradient 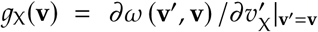 for the corresponding virulence, until **v** = **v**\* is attained, with *g*_J_(**v**\*) = *g*_A_(**v**\*) = 0. For more detail, see Appendix (A5).

We investigate the effects of two stage-structure characteristics: (i) post-maturation span *θ*_A_ and (ii) stage-assortativity *ρ* on the evolutionary outcomes (i.e., CSS; Appendix A6-9). We use the following default parameter values: *r* = 6, *κ* = 0.06, *m*_J_ = 1, *α*_J_ = *α*_A_ = 1, *k*_J_ = *k*_A_ = 1, *b*_J_ = *b*_A_ = 10, while varying *m*_A_, *u* (thus *θ*_A_) and *ρ*. That is, the parameter values are symmetric for juveniles and adults. We subsequently check the effects of the differences in *α* (susceptibility), *k* (efficiency of exploitation for transmission), and *b* (upper bound in infectiousness). We also check whether recovery, tolerance, or fecundity shift in the hosts can affect the results. In addition, we examined the outcomes when we assume density-dependent rather than frequency dependent transmission in the dynamics. Finally, we investigated more general transmission pathways with (i) *σ*_JJ_, *σ*_AA_, and *σ*_JA_ = *σ*_AJ_ all varying freely, and (ii) *σ*_JJ_ = 1 – *σ*_AJ_ and *σ*_AA_ = 1 – *σ*_JA_ both varying freely.

## Results

The selection gradient for adult-virulence reads:

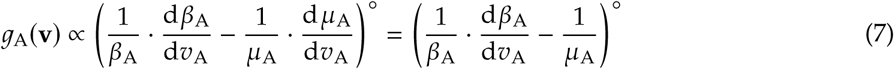

(where º represents neutrality, **v**′ = **v**; Appendix A5), which is consistent with a number of previous studies: under the transmission-virulence trade-off, higher exploitation is expected to increase the infectiousness (i.e., a marginal benefit) at the immediate (marginal) cost owing to a reduced infectious period (Day 2001; Gandon *et al.* 2001; Day & Proulx 2004; Gandon 2004; Alizon *et al.* 2009; Cressler *et al.* 2016). This is because the reproductive success of parasites infecting adults is, in effect, determined by a single transmission pathway, from adults to any susceptible hosts in the population, regardless of the stage structure. Therefore, the direction of selection on adult virulence is completely determined by the balance between such marginal benefits and costs, regardless of the characteristics of hosts’ stage-structure. The resulting CSS for adult-virulence is always 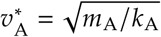 which is independent of any demographic and disease characteristics of juveniles. We can immediately see that the evolutionary outcome of adult-virulence increases with adult natural mortality (reviewed in Alizon *et al.* 2009; Cressler *et al.* 2016).

In contrast, juvenile-virulence in the model is influenced by additional costs associated with hosts’ stage structure. This is because the parasites infecting the juveniles can spread either from the juvenile (to any susceptible hosts), or from the adult who has successfully matured from the juvenile stage. To ease biological interpretation, we present the reproductive value based form of the selection gradient (Fisher 1958; Taylor 1990; Caswell 2001; Frank 1998; Grafen 2006; Lion 2018). Reproductive values give a proper weighting of fitness effects for age-classes, by taking the contributions of classes to future gene pool into account (Fisher 1958; Taylor 1990; Caswell 2001; Frank 1998; Grafen 2006; Lion 2018). Using reproductive values, *g*_J_(**v**) reads:

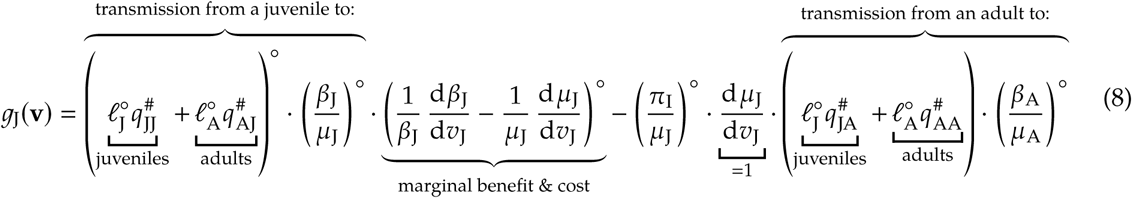

(Appendix A6-8), where 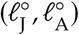 represents the pair of individual reproductive values of the parasites infecting juvenile and adult hosts (defined by the left eigenvector of **G** at neutrality; Appendix A6) and the factor 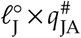 for instance represents the reproductive success due to transmission from an adult host to a juvenile per propagule-production. The first term is multiplied by 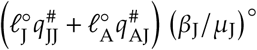, which represents the reproductive success of a parasite infecting juveniles, who can receive the marginal benefit due to increased infectiousness 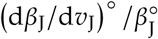 but pay the marginal cost due to the reduction in infectious periods 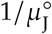 as in the selection gradient for adult-virulence (Eqn (7)). In addition, juvenile-virulence incurs the additional cost associated with the reduced maturation probability (π_I_/*µ*_J_)º and the subsequent loss of expected reproductive success via adult hosts that the parasites could otherwise gain through the maturation of the juvenile hosts, 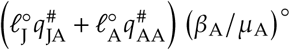 (the reproductive success of a parasite infecting adults). Hence, Eqn (8) clearly captures the selection forces on juvenile-virulence, including the marginal benefits, marginal costs, and maturation-mediated costs. The expression for the CSS of juvenile-virulence, 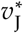, is analytically intractable, and as such we numerically evaluated *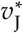* by jointly solving the wild-type ODEs in Eqn (1) and the selection gradients in Eqns (7) and (8).

To assess when selection favors higher juvenile-virulence than adult-virulence, we quantified 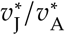 as a function of the assortatvity (*ρ*, abscissa) and post-maturation span (*θ*_A_, ordinate; Fig 3). We found that either disassortative hosts with a long post-maturation span or assortative hosts with a short post-maturation span select for higher virulence against juveniles. This result slightly changes given stage-specific mortality rates (*m*_J_ ≠ *m*_A_) such that a higher mortality for juveniles can bias the outcomes towards higher virulence for juveniles, but the general trend is robust (Fig 3A-C). Also, the combination of disassortativity and long post-maturation span (one of the conditions favoring higher virulence against juveniles) leads to parasite extinction as a result of overexploitation against juveniles (Fig 3B, C; see Appendix A10).

By relaxing the assumptions of the symmetry in disease-related parameters *k*_J_, *k*_A_ (efficiency of exploitation for transmission), *b*_J_, *b*_A_ (maximum infectiousness), and *α*_J_, *α*_A_ (susceptibility) for juveniles and adults, or by incorporating recovery or tolerance, we showed that the results are robust and qualitatively unchanged (Appendix B). In addition, we showed that density-dependent transmission yields quantitatively similar results (Appendix B). We further found that fecundity shift (i.e., infected adults have different fecundity output than do susceptible hosts, as in *Daphnia*; Hite *et al.* 2017) have minor impacts upon the results. Finally, we found that varying *σ*_JJ_, *σ*_AJ_, and *σ*_AA_ yields the quantitatively similar results (Appendix C). Therefore, we conclude that the combined effects of maturation and assortativity are critical to the evolution of virulence.

## Discussion

We have shown how parasites may be subject to different selective pressures when they infect juveniles as opposed to adults. Our key insight is that the combination between maturation rates and transmission pathway determines the evolutionary outcomes of juvenile-virulence. Higher virulence against juveniles is favored either if: (i) the adult-stage is relatively long and the transmission pathway is disassortative (between age-class transmission likely; Fig 3, left top zone), or (ii) the juvenile-stage is relatively long and the transmission pathway is assortative (transmission occurs preferentially within classes; Fig 3, right bottom zone). These results can be understood as follows: when the post-maturation span is long and the transmission pathway is disassortative, adult hosts are abundant in the population and the transmission from juveniles to adults is more likely than between juveniles. The higher availability of adult hosts selects for higher transmission and therefore virulence in juveniles in order to exploit more abundant resource of adults. Equivalent reasoning explains higher juvenile-virulence when maturation is fast and hosts are assortative. As such the hosts’ demography alongside the maturation of juveniles strongly affects the evolutionary outcomes of parasite virulence. Spatial and/or temporal segregation in the niches of juveniles and adults therefore has the potential to drive the evolution of differential virulence. Our novel result is therefore that virulence is highly sensitive to stage-structured life history characteristics of hosts including ontogeny and any associated, spatiotemporal niche shifts.

Higher parasite exploitation against juveniles incurs an additional cost associated with increased maturation failure. In contrast, the evolutionary outcome of adult virulence is fully understood from the classic perspective of optimizing the secondary infections from infected adults. Therefore, sources of heterogeneity in hosts can lead to different predictions than classic virulence evolution theory based on the marginal value theorem, as claimed in a recent conceptual review (Lion & Metz 2018). Our novel results arise because we explicitly assumed stage-structure with maturation from juveniles to adults and reproduction by adults, rather than more generic host heterogeneity, e.g., multiple host species (Regoes *et al.* 2000; Gandon 2004; Osnas & Dobson 2011), vaccination (Gandon *et al.* 2001; Gandon *et al.* 2003; Yates *et al.* 2006; Zurita-Gutiérrez & Lion 2015), or sex (Úbeda & Jansen 2016).

Finding examples of stage-specific virulence in empirical systems can be difficult due to the intricacies of specific host-pathogen systems. Stage-related trends in virulence can be complicated by age-related trends in maternal immunity, adaptive immunity, and exposure rate, alongside the impact of maladaptation and immunopathogenicity (Hudson & Dobson 1995; Wilson *et al.* 2002). Additionally, studies looking at age-related virulence or case mortality do not exclusively look at differences between adult and juvenile stages and may focus on old age-mediated declines in immunocompetence. However, despite these issues, we found data on several empirical systems in an intensive literature review (Appendix D) where age-biased virulence effects can be distinguished from the other factors and which lend support to our predictions and offer opportunities for testing our hypotheses (Fig 1; also see Appendix D). For most of these systems, we were unable to find data on the assortativity of transmission which therefore limited our ability to make conclusions about trends in the data. However, both of the two wildlife systems for which we found data describing all three of our variables (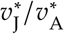, post-maturation lifespan, and transmission assortativity) matched our model’s predictions (Fig 1). Wanelik *et al.* (2017) showed that Great Island Virus (GIV) transmission in guillemots (*Uria aalge*) is assortative across age classes because of the spatial structure of breeding grounds. GIV is transmitted by poorly motile ticks and prebreeding stages of guillemots do not enter breeding areas of the colony. As a consequence, the virus does not readily transmit between guillemot age-stages (Wanelik *et al.* 2017). Previous work on guillemot life history shows that the birds spend more than three quarters of their lifespan as mature breeders (Harris & Wanless 1995), and therefore the combination of assortative transmission and fast maturation predicts that GIV should be more virulent in breeders. In line with the predictions of our model, infection associated mortality risk is 0.63 times lower for juveniles than for adults (Nunn *et al.* 2006).

**Figure 1:**
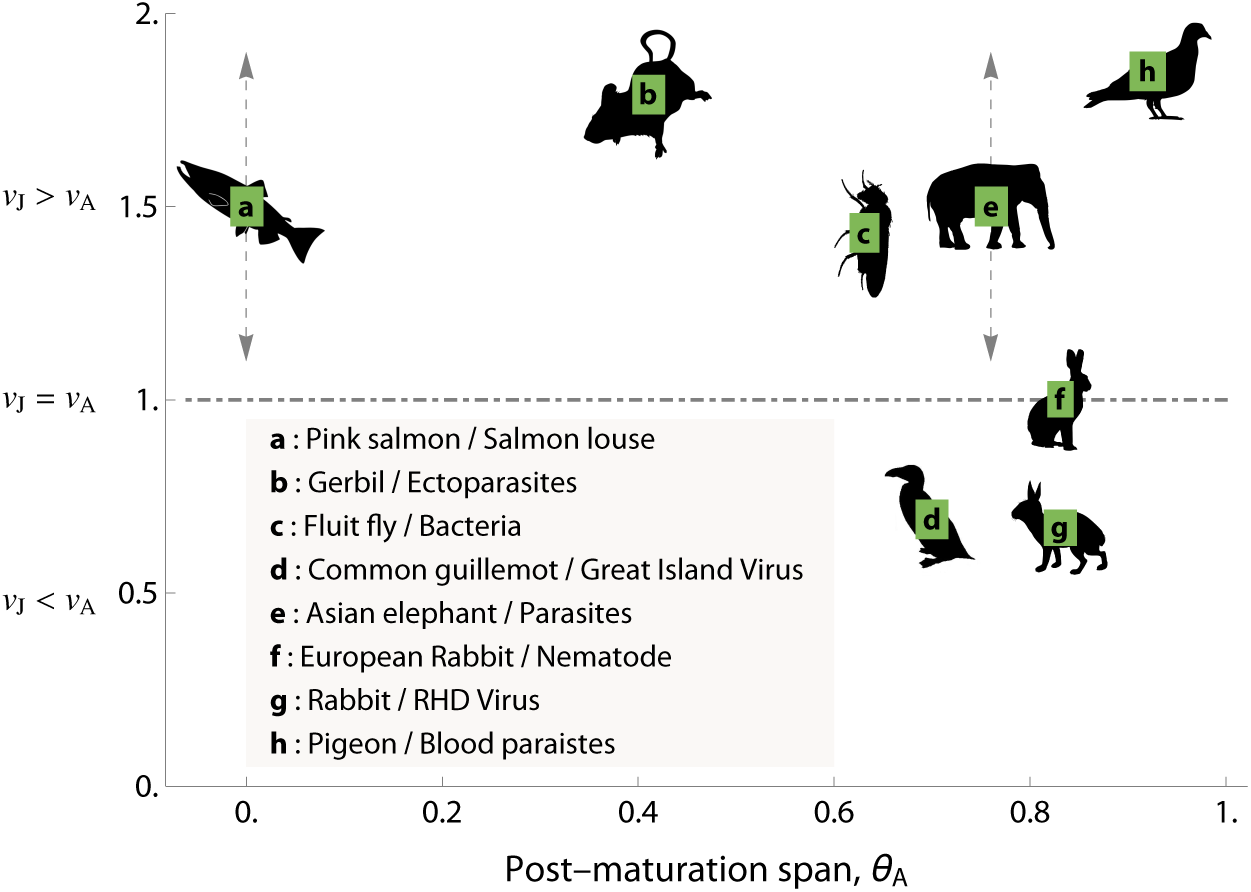
A graphical representation of the empirical data on stage-specific virulence. Each position of labels (from a to h, each within a green square) corresponds to a reported value of *v*_J_/ *v*_A_. In the following, H indicates “host” while P indicates “parasite(s)/pathogen(s)”. (a) H: Pink Salmon (*Oncorhynchus gorbuscha*); P: Salmon Louse (Heard 1991; Jones *et al.* 2008). (b) H: Gerbil (*Gerbillus andersoni*); P: Ectoparasites, including fleas (*Synosternus cleopatrae*), mites (*Androlaelaps centrocarpus, A. insculptus, A. hirst i, A. marshalli* and *A. androgynus*), and ticks (*Hyalomma impeltatum*) (Wassif & Soliman 1980; Delany 1986; Hawlena *et al.* 2006). (c) H: Fluit fly (*Drosophila melanogaster*) P: Bacteria (*Pseudomonas entomophila*) (Vodovar *et al.* 2005; Luckinbill *et al.* 1984); (d) H: Common guillemot (*Uria aalge*); P: Great Island Virus (Harris & Wanless 1995; Nunn *et al.* 2006; Wanelik *et al.* 2017). (e) H: Asian elephant (*Elephas maximus*); P: Parasites (Sukumar *et al.* 1997; Lynsdale *et al.* 2017). (f) H: European rabbit (*Oryctolagus cuniculus*); P: Nematode (von Holst *et al.* 2002; Cornell *et al.* 2008). (g) H: Rabbits (*Leporidae*); P: RHD Virus (Morisse *et al.* 1991; Reluga *et al.* 2007). (h) H: Pigeon (*Columba livia*); P: Blood parasites (Lack 1968; Holmes & Ottinger 2003; Sol *et al.* 2003). For (a) and (e), we were able to find qualitative data (*v*_J_ > *v*_A_), but not quantitative ones. As such, we placed them at the height of 1.5 (>1), and indicated variations by dashed, gray arrows thereon. Data behind this figure is shown in SI Table in Appendix D. Animal drawings are from phylopic.org, with full credit in Appendix D.

**Figure 2:**
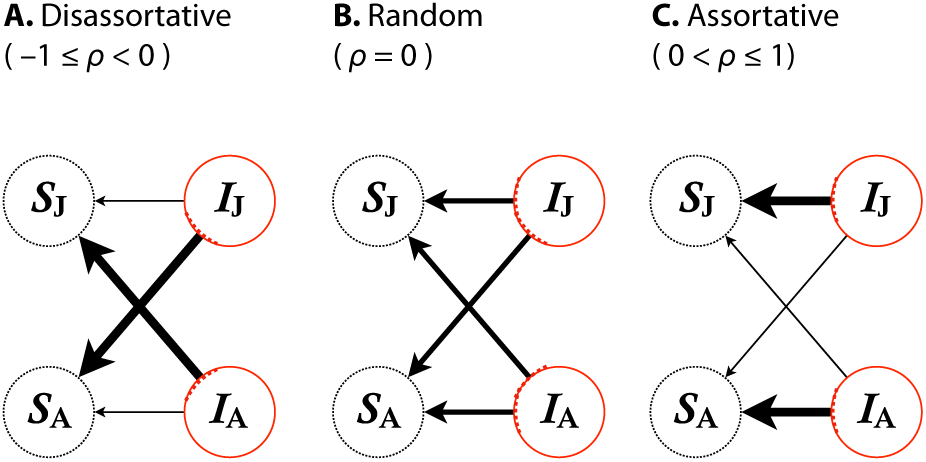
A schematic illustration of the assortativity parameter *ρ*. Negative assortativity indicates that contacts occur more frequently between stages than within stages (panel A). The transmission pathway is unbiased (random) when *ρ* = 0 (panel B). Positive assortativity indicates that transmission occurs more frequently within stages than between stages (panel C).

**Figure 3:**
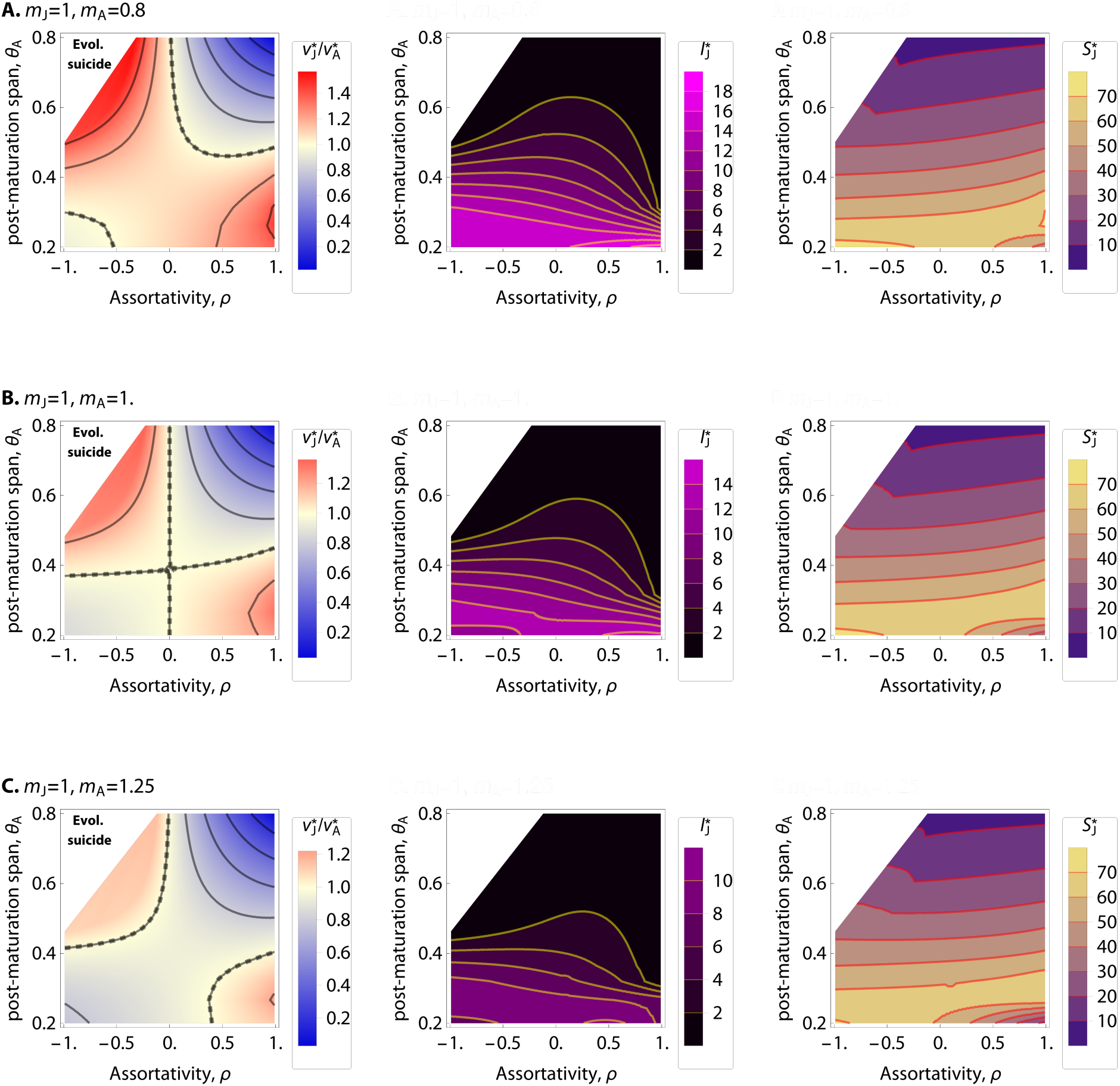
Left panels: Evolutionary outcomes of relative virulence 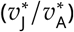, in which red color indicates 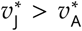 and blue color indicates 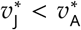. Color scales used are the same in the three panels. Middle panels: Densities of infected juveniles at equilibrium, 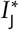. Right panels: Densities of susceptible juveniles at equilibrium, 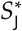. In each panel, abscissa: assortativity; ordinate: post-maturation span *θ*_A_; from (A) to (C): *m*_A_ = 0.8, 1.0, 1.25 as indicated; white zone: evolutionary suicide; dotted curve: 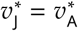 (equal virulence); parameters: default values. Middle and right panel can clearly demonstrate that evolutionary suicide does occur in the white zone, as 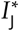 is very small but 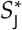 is not.

In the second example, Jones *et al.* (2008) showed that salmon louse caused morality in juvenile pink salmon (*Oncorhynchus gorbuscha*), but had no effect on mortality risk for adults. Salmon louse is also assortatively transmitted between age classes, because pink salmon have strict two year lifespans where they are only ever associated with individuals of their same age class (Heard 1991; Krkošek *et al.* 2007). The salmon only reproduce once at the very end of their lives (semelparity), and therefore have a short adult period. This short post-maturation stage and assortative transmission correctly predicts the higher salmon louse virulence in juveniles.

Better data on mixing matrices for more disease systems would provide interesting insights into the maintenance of either high juvenile or high adult virulence. One system where these insights could prove especially important is in Bd (*Batrachochytrium dendrobatidis*, or chytrid fungus) infection in frogs, which has been causing catastrophic worldwide declines in frog populations (Kilpatrick *et al.* 2010). Bd infection has been shown to have different virulence effects in the different frog life stages (Briggs *et al.* 2010; Medina *et al.* 2015; Hite *et al.* 2016) and these effects also vary by frog species (Berger *et al.* 1998; Blaustein *et al.* 2005). Recent work has shown that adult virulence in several frog populations has not decreased even after 20 years of Bd presence (Voyles *et al.* 2018). Already, frog demography has been implicated as an important factor for population persistence in the face of Bd with frog species where adults move away from breeding waters being more resistant to population declines (Lips *et al.* 2006; McCaffery *et al.* 2015), and frogs in habitats with multi-year larvae having more severe epidemics because the older stages maintain high levels of infection that then spill over to infect other stages and species (Medina *et al.* 2015; Hite *et al.* 2016). Changes in the assortativity of mixing clearly has important implications for disease transmission across stages, and our model suggests that it could also have implications for the maintenance of high virulence in different age stages.

While data on age-related transmission pathways are difficult to find in wildlife populations, a wealth of mixing data exists for humans (Mossong *et al.* 2008; Rohani *et al.* 2010). These suggest that contacts relevant for the transmission of directly transmitted pathogens are highly assortative by age. Given that humans have a long juvenile period in the context of our model (Bogin & Smith 1996), our model predicts that directly transmitted human-adapted pathogens would be selected to have higher virulence in naive adults, but cannot be applied to pathogens that are adapted to multiple host species or are vector- or sexually transmitted. Measles and chickenpox, both being directly transmitted human specific pathogens, have case mortality rates that are 1.7 times and 23–29 times (respectively) higher in unvaccinated, unexposed adults than in juveniles (Orenstein *et al.* 2004; Heininger & Seward 2006). Poliovirus also has a 8.75 times higher case mortality rate for adults than juveniles (Weinstein *et al.* 1952) though the assortativity of its transmission seems to depend on the relative contribution of direct (oral-oral) and environmental (fecal-oral) transmission (Blake *et al.* 2014). While the evolutionary drivers of human pathogens are often complicated, we posit that chickenpox (varicella virus) virulence in humans proves an intriguing case study. The higher mortality risk in adults corresponds to increased viral titers with age, suggesting that there may be a tradeoff between virulence and transmission (Malavige *et al.* 2008). Perhaps most interestingly, while varicella virus infects many cell types, T cell infection is thought to be important for transport and pathogenesis (Zerboni & Arvin 2016). Therefore, age-related trends in T-cell abundance could be implicated in chickenpox pathogenesis, although this relationship is complicated by the fact that VSV-specific T-cell responses are also correlated with decreased viral titer and diminish with age (Erkeller-Yuksel *et al.* 1992; Nader *et al.* 1995; Malavige *et al.* 2008). Altogether, the implications for human pathogens entail deeper understanding of mechanisms for immunocompetence and modes of transmission (sexual and/or environmental), but the examples point towards one mechanism that may contribute to the mediation of age-specific virulence in human pathogens.

Our models have implications for disease management especially in farmed and other managed animal populations. For instance, if the post-maturation span is short (i.e. if *u* is small), then artificial restriction of the physical contacts between stages is predicted to select for higher virulence. However, if the post-maturation span is long, restricting the contacts into juvenile-juvenile and adult-adult (by e.g., separating the cohorts) can lead to the parasite extinction as a result of overexploitation against the juveniles. These contrasting outcomes can occur for any given host species, depending on how management modulates host stage-structure. Therefore to prevent evolutionary changes towards higher virulence, we should carefully take into account the cohort structure.

For simplicity and tractability, we chose to use simple two stage models rather than continuous “infection-age” models. Future studies that capture more continuous age structure are an important next step. Also, although we assumed that parasites can express conditional virulence depending on the stage of the hosts they infect with, more data are needed to test this idea. In addition, coevolutionary models and multiple infections are both likely to give further important insights to the determinants of age-dependent disease interactions in nature. Our approach offers the basis for modeling these coevolutionary dynamics between hosts and parasites when there is stage structure.

## Acknowledgement

We thank two anonymous reviewers for their helpful comments, and Virpi Lummaa and Carly Lynsdale for sharing the data on asian elephant demography. This study is supported by the National Science Foundation Graduate Research Fellowship Program (1752814) and The Natural Environment Research Council (NERC; NE/K014617/1).

Author contribution: All authors conceived and designed the study, RI carried out model analyses, RI and EV drafted the initial version of the manuscript, EV surveyed empirical literature, and all authors contributed to later versions of the manuscript.

