## Appendix for "The evolution of stage-specific virulence: differential selection of parasites in juveniles"

January 28, 2019

**C Generalized pathway structure** **18**

**D Empirical data figure and credits**

### A Invasion analysis

#### A.1 Generalized ODE

The epidemiological dynamics is given by:

$$\begin{aligned}
\frac{\mathrm{d}S_\mathrm{J}}{\mathrm{d}t} &= \left(r - \kappa(S_\mathrm{A} + I_\mathrm{A})\right) \cdot (S_\mathrm{A} + I_\mathrm{A}) - \left(u + \phi_\mathrm{JA} + \phi_\mathrm{JJ} + m_\mathrm{J}\right) S_\mathrm{J} + \gamma_\mathrm{J} I_\mathrm{J}, \\
\frac{\mathrm{d}S_\mathrm{A}}{\mathrm{d}t} &= u S_\mathrm{J} - \left(m_\mathrm{A} + \phi_\mathrm{AJ} + \phi_\mathrm{AA}\right) S_\mathrm{A} + \gamma_\mathrm{A} I_\mathrm{A}, \\
\frac{\mathrm{d}I_\mathrm{J}}{\mathrm{d}t} &= \left(\phi_\mathrm{JA} + \phi_\mathrm{JJ}\right) S_\mathrm{J} - \left(u + m_\mathrm{J} + v_\mathrm{J} + \gamma_\mathrm{J}\right) I_\mathrm{J}, \\
\frac{\mathrm{d}I_\mathrm{A}}{\mathrm{d}t} &= \left(\phi_\mathrm{AJ} + \phi_\mathrm{AA}\right) S_\mathrm{A} + u I_\mathrm{J} - \left(m_\mathrm{A} + v_\mathrm{A} + \gamma_\mathrm{A}\right) I_\mathrm{A},
\end{aligned} \tag{A.1}$$

with the notation explained in the main text; here, for the sake of generality, we incorporated recovery $\gamma_\mathrm{J}, \gamma_\mathrm{A}$, which we will use later. Solving the system gives two equilibria: one is disease free $(S_\mathrm{J}^{(0)}, S_\mathrm{A}^{(0)}, 0, 0)$, and the other is endemic $(S_\mathrm{J}^{\#}, S_\mathrm{A}^{\#}, I_\mathrm{J}^{\#}, I_\mathrm{A}^{\#})$.

#### A.2 Stage-period

In this subsection, we will restrict our attention to the disease-free subsystem:

$$\begin{aligned}
\frac{\mathrm{d}S_\mathrm{J}}{\mathrm{d}t} &= (r - \kappa S_\mathrm{A}) S_\mathrm{A} - \left(u + m_\mathrm{J}\right) S_\mathrm{J}, \\
\frac{\mathrm{d}S_\mathrm{A}}{\mathrm{d}t} &= u S_\mathrm{J} - m_\mathrm{A} S_\mathrm{A}.
\end{aligned} \tag{A.2}$$

First, the probability of successful maturation is given by:

$$\pi_\mathrm{S} = \frac{u}{u + m_\mathrm{J}}. \tag{A.3}$$

Second, consider two random variables: the duration of time a host individual spends as a juvenile, denoted $T_\mathrm{J}$, and the duration of time a host individual spends as an adult, denoted $T_\mathrm{A}$. The fate of a juvenile is (i) to die as a juvenile or (ii) to successfully mature and die as an adult. For the former case, which occurs with probability $1 - \pi_\mathrm{S}$, the random variable $T_\mathrm{J}$ follows an exponential distribution with mean $1/(u + m_\mathrm{J})$ while $T_\mathrm{A} \equiv 0$. With probability $\pi_\mathrm{S}$, the latter happens, in which case, the bivariate random variables $(T_\mathrm{J}, T_\mathrm{A})$ follow the two dimensional exponential distribution, given by:

$$\left(T_\mathrm{J}, T_\mathrm{A}\right) \text{ follows } \left(u + m_\mathrm{J}\right) e^{-(u+m_\mathrm{J})T_\mathrm{J}} \cdot m_\mathrm{A} e^{-m_\mathrm{A} T_\mathrm{A}}. \tag{A.4}$$

Therefore, the expectation of $T_\mathrm{A}/(T_\mathrm{J}+T_\mathrm{A})$ is given by:

$$\theta_\mathrm{A} = (1-\pi_\mathrm{S})\cdot 0 + \pi_\mathrm{S}\cdot \iint_0^\infty \frac{T_\mathrm{A}}{T_\mathrm{J}+T_\mathrm{A}} \left(u+m_\mathrm{J}\right) e^{-\left(u+m_\mathrm{J}\right)T_\mathrm{J}} \cdot m_\mathrm{A} e^{-m_\mathrm{A}T_\mathrm{A}} \,\mathrm{d}T_\mathrm{J}\,\mathrm{d}T_\mathrm{A}. \tag{A.5}$$

To calculate the integral, we carry out the variable transformation by:

$$L := T_\mathrm{J}+T_\mathrm{A}, f_\mathrm{A} := \frac{T_\mathrm{A}}{T_\mathrm{J}+T_\mathrm{A}} \Longleftrightarrow T_\mathrm{J} = L(1-f_\mathrm{A}),\ T_\mathrm{A} = Lf_\mathrm{A}, \tag{A.6}$$

with the corresponding Jacobian of the variable transformation:

$$\frac{\partial\left(T_\mathrm{J},T_\mathrm{A}\right)}{\partial\left(L,f_\mathrm{A}\right)} := \left|\det\begin{pmatrix} \frac{\partial T_\mathrm{J}}{\partial L} & \frac{\partial T_\mathrm{J}}{\partial f_\mathrm{A}} \\ \frac{\partial T_\mathrm{A}}{\partial L} & \frac{\partial T_\mathrm{A}}{\partial f_\mathrm{A}} \end{pmatrix}\right| = L\,(>0)\,. \tag{A.7}$$

Noting that $0 \le f_\mathrm{A} \le 1$, we have:

$$\begin{aligned} \theta_\mathrm{A} &= \frac{u}{u+m_\mathrm{J}} \iint_0^\infty \frac{T_\mathrm{A}}{T_\mathrm{J}+T_\mathrm{A}} \left(u+m_\mathrm{J}\right) e^{-\left(u+m_\mathrm{J}\right)T_\mathrm{J}} \cdot m_\mathrm{A} e^{-m_\mathrm{A}T_\mathrm{A}} \,\mathrm{d}T_\mathrm{J}\,\mathrm{d}T_\mathrm{A} \\ &= \frac{u}{u+m_\mathrm{J}} \int_0^1 \int_0^\infty f_\mathrm{A}\cdot\left(u+m_\mathrm{J}\right)\cdot m_\mathrm{A}\cdot e^{-\left((u+m_\mathrm{J})(1-f_\mathrm{A})+m_\mathrm{A}f_\mathrm{A}\right)L} L\,\mathrm{d}L\,\mathrm{d}f_\mathrm{A}. \end{aligned} \tag{A.8}$$

Integrating with respect to $L$ firstly and then integrating with respect to $f_\mathrm{A}$, we have:

$$\theta_\mathrm{A} = \frac{u}{u+m_\mathrm{J}-m_\mathrm{A}} \left(1 + \frac{m_\mathrm{A}}{u+m_\mathrm{J}-m_\mathrm{A}} \cdot \log\left(\frac{m_\mathrm{A}}{u+m_\mathrm{J}}\right)\right) \tag{A.9}$$

as shown in the main text.

Note that if $u + m_\mathrm{J} = m_\mathrm{A}$, then $\theta_\mathrm{A}$ is of the form "0/0". As such $\theta_\mathrm{A}$ is interpreted as the limit $\lim_{m_\mathrm{A}\to u+m_\mathrm{J}} = \pi_\mathrm{S}/2$, which is the probability of maturation ($\pi_\mathrm{S}$) times the conditional expectation of the fraction of sub-lifespan as an adult (given that a sampled adult host has matured into an adult). This calculation is obtained by setting $\exp(\varepsilon) := m_\mathrm{A}/(u+m_\mathrm{J})$ and using the Taylor expansion $\exp(\varepsilon) = 1 + \varepsilon + \frac{\varepsilon^2}{2} + O(\varepsilon^3)$ where $O()$ represents the Landau's big-$O$ for $\varepsilon \to +0$. Exact computation including the evaluation of integral is shown in a Mathematica-code (SI Fig 1).

#### A.3 Mutant dynamics

Hereafter, without special remarks, we will assume that $\rho \lesssim 1$ (i.e., transmission can occur between classes). When $\rho = 1$, as shown in Osnas & Dobson (2011), a special treatment is needed.

Evaluating the stage-period requires variable-transformation,
but Mathematica can skip this task.

```
In[•]:= Assuming[mA > 0 && mJ > 0 && u > 0, u / (u + mJ) *
          Integrate[tA / (tA + tJ) * (u + mJ) * mA *
            Exp[- (u + mJ) * tJ] * Exp[- (mA) * tA],
           {tJ, 0, +∞}, {tA, 0, +∞}]];

In[•]:= % - u / (u + mJ - mA) *
          (1 + mA / (u + mJ - mA) * Log[mA / (u + mJ)]) //
         Simplify

Out[•]= 0
```

...as desired.

```
In[•]:= Limit[%%, mA → +u + mJ]

Out[•]= u / (2 (mJ + u))
```

...as desired.

SI Figure 1: Mathematica code for evaluating the stage-period.

The dynamics governing the mutant's growth rate (mutant dynamics) reads:

$$\begin{aligned}
\frac{\mathrm{d}I'_{\mathrm{J}}}{\mathrm{d}t} &= \left(\phi'_{\mathrm{JA}} + \phi'_{\mathrm{JJ}}\right) S^{\#}_{\mathrm{J}} - \left(u + m_{\mathrm{J}} + v'_{\mathrm{J}} + \gamma_{\mathrm{J}}\right) I'_{\mathrm{J}} \\
&= \left(\phi'_{\mathrm{JA}} + \phi'_{\mathrm{JJ}}\right) S^{\#}_{\mathrm{J}} - \mu'_{\mathrm{J}} I'_{\mathrm{J}}, \\
\frac{\mathrm{d}I'_{\mathrm{A}}}{\mathrm{d}t} &= \left(\phi'_{\mathrm{AJ}} + \phi'_{\mathrm{AA}}\right) S^{\#}_{\mathrm{A}} + u I'_{\mathrm{J}} - \left(m_{\mathrm{A}} + v'_{\mathrm{A}} + \gamma_{\mathrm{A}}\right) I'_{\mathrm{A}} \\
&= \left(\phi'_{\mathrm{AJ}} + \phi'_{\mathrm{AA}}\right) S^{\#}_{\mathrm{A}} + u I'_{\mathrm{J}} - \mu'_{\mathrm{A}} I'_{\mathrm{A}}.
\end{aligned} \tag{A.10}$$

Here,

$$\begin{aligned}
\phi'_{\mathrm{JJ}} &= \frac{\alpha_{\mathrm{J}} \sigma_{\mathrm{JJ}} \beta'_{\mathrm{J}} I'_{\mathrm{J}}}{S^{\#}_{\mathrm{J}} + S^{\#}_{\mathrm{A}} + I^{\#}_{\mathrm{J}} + I^{\#}_{\mathrm{A}}}, \\
\phi'_{\mathrm{JA}} &= \frac{\alpha_{\mathrm{J}} \sigma_{\mathrm{JA}} \beta'_{\mathrm{A}} I'_{\mathrm{A}}}{S^{\#}_{\mathrm{J}} + S^{\#}_{\mathrm{A}} + I^{\#}_{\mathrm{J}} + I^{\#}_{\mathrm{A}}}, \\
\phi'_{\mathrm{AJ}} &= \frac{\alpha_{\mathrm{A}} \sigma_{\mathrm{AJ}} \beta'_{\mathrm{J}} I'_{\mathrm{J}}}{S^{\#}_{\mathrm{J}} + S^{\#}_{\mathrm{A}} + I^{\#}_{\mathrm{J}} + I^{\#}_{\mathrm{A}}}, \\
\phi'_{\mathrm{AA}} &= \frac{\alpha_{\mathrm{A}} \sigma_{\mathrm{AA}} \beta'_{\mathrm{A}} I'_{\mathrm{A}}}{S^{\#}_{\mathrm{J}} + S^{\#}_{\mathrm{A}} + I^{\#}_{\mathrm{J}} + I^{\#}_{\mathrm{A}}}.
\end{aligned} \tag{A.11}$$

#### A.4 Invasion fitness and invadability condition

Linearizing the mutant dynamics around the endemic equilibrium, we get a corresponding Jacobian:

$$\begin{aligned}
\mathbf{J}' &= \begin{pmatrix} \dfrac{\alpha_{\mathrm{J}} S^{\#}_{\mathrm{J}} \sigma_{\mathrm{JJ}} \beta'_{\mathrm{J}}}{S^{\#}_{\mathrm{J}} + S^{\#}_{\mathrm{A}} + I^{\#}_{\mathrm{J}} + I^{\#}_{\mathrm{A}}} & \dfrac{\alpha_{\mathrm{J}} S^{\#}_{\mathrm{J}} \sigma_{\mathrm{JA}} \beta'_{\mathrm{A}}}{S^{\#}_{\mathrm{J}} + S^{\#}_{\mathrm{A}} + I^{\#}_{\mathrm{J}} + I^{\#}_{\mathrm{A}}} \\ \dfrac{\alpha_{\mathrm{A}} S^{\#}_{\mathrm{A}} \sigma_{\mathrm{AJ}} \beta'_{\mathrm{J}}}{S^{\#}_{\mathrm{J}} + S^{\#}_{\mathrm{A}} + I^{\#}_{\mathrm{J}} + I^{\#}_{\mathrm{A}}} & \dfrac{\alpha_{\mathrm{A}} S^{\#}_{\mathrm{A}} \sigma_{\mathrm{AA}} \beta'_{\mathrm{A}}}{S^{\#}_{\mathrm{J}} + S^{\#}_{\mathrm{A}} + I^{\#}_{\mathrm{J}} + I^{\#}_{\mathrm{A}}} \end{pmatrix} - \begin{pmatrix} \mu'_{\mathrm{J}} & 0 \\ -u & \mu'_{\mathrm{A}} \end{pmatrix} \\
&= \mathbf{B}' - \mathbf{D}'.
\end{aligned} \tag{A.12}$$

The next generation matrix $\mathbf{G}'$ is given by:

$$\begin{aligned}
\mathbf{G}' &= \mathbf{B}'\left(\mathbf{D}'\right)^{-1} \\
&= \begin{pmatrix} a'_{\mathrm{JJ}} & a'_{\mathrm{JA}} \\ a'_{\mathrm{AJ}} & a'_{\mathrm{AA}} \end{pmatrix} \\
&= \begin{pmatrix} \dfrac{\alpha_{\mathrm{J}} S^{\#}_{\mathrm{J}} \sigma_{\mathrm{JJ}} \beta'_{\mathrm{J}}}{S^{\#}_{\mathrm{J}} + S^{\#}_{\mathrm{A}} + I^{\#}_{\mathrm{J}} + I^{\#}_{\mathrm{A}}} & \dfrac{\alpha_{\mathrm{J}} S^{\#}_{\mathrm{J}} \sigma_{\mathrm{JA}} \beta'_{\mathrm{A}}}{S^{\#}_{\mathrm{J}} + S^{\#}_{\mathrm{A}} + I^{\#}_{\mathrm{J}} + I^{\#}_{\mathrm{A}}} \\ \dfrac{\alpha_{\mathrm{A}} S^{\#}_{\mathrm{A}} \sigma_{\mathrm{AJ}} \beta'_{\mathrm{J}}}{S^{\#}_{\mathrm{J}} + S^{\#}_{\mathrm{A}} + I^{\#}_{\mathrm{J}} + I^{\#}_{\mathrm{A}}} & \dfrac{\alpha_{\mathrm{A}} S^{\#}_{\mathrm{A}} \sigma_{\mathrm{AA}} \beta'_{\mathrm{A}}}{S^{\#}_{\mathrm{J}} + S^{\#}_{\mathrm{A}} + I^{\#}_{\mathrm{J}} + I^{\#}_{\mathrm{A}}} \end{pmatrix} \begin{pmatrix} \dfrac{1}{\mu'_{\mathrm{J}}} & 0 \\ \dfrac{u}{\mu'_{\mathrm{J}}\mu'_{\mathrm{A}}} & \dfrac{1}{\mu'_{\mathrm{A}}} \end{pmatrix} \\
&= \begin{pmatrix} \dfrac{\alpha_{\mathrm{J}} S^{\#}_{\mathrm{J}} \sigma_{\mathrm{JJ}} \beta'_{\mathrm{J}}}{S^{\#}_{\mathrm{J}} + S^{\#}_{\mathrm{A}} + I^{\#}_{\mathrm{J}} + I^{\#}_{\mathrm{A}}} \cdot \dfrac{1}{\mu'_{\mathrm{J}}} + \dfrac{\alpha_{\mathrm{J}} S^{\#}_{\mathrm{J}} \sigma_{\mathrm{JA}} \beta'_{\mathrm{A}}}{S^{\#}_{\mathrm{J}} + S^{\#}_{\mathrm{A}} + I^{\#}_{\mathrm{J}} + I^{\#}_{\mathrm{A}}} \cdot \dfrac{u}{\mu'_{\mathrm{J}}\mu'_{\mathrm{A}}} & \dfrac{\alpha_{\mathrm{J}} S^{\#}_{\mathrm{J}} \sigma_{\mathrm{JA}} \beta'_{\mathrm{A}}}{S^{\#}_{\mathrm{J}} + S^{\#}_{\mathrm{A}} + I^{\#}_{\mathrm{J}} + I^{\#}_{\mathrm{A}}} \cdot \dfrac{1}{\mu'_{\mathrm{A}}} \\ \dfrac{\alpha_{\mathrm{A}} S^{\#}_{\mathrm{A}} \sigma_{\mathrm{AJ}} \beta'_{\mathrm{J}}}{S^{\#}_{\mathrm{J}} + S^{\#}_{\mathrm{A}} + I^{\#}_{\mathrm{J}} + I^{\#}_{\mathrm{A}}} \cdot \dfrac{1}{\mu'_{\mathrm{J}}} + \dfrac{\alpha_{\mathrm{A}} S^{\#}_{\mathrm{A}} \sigma_{\mathrm{AA}} \beta'_{\mathrm{A}}}{S^{\#}_{\mathrm{J}} + S^{\#}_{\mathrm{A}} + I^{\#}_{\mathrm{J}} + I^{\#}_{\mathrm{A}}} \cdot \dfrac{u}{\mu'_{\mathrm{J}}\mu'_{\mathrm{A}}} & \dfrac{\alpha_{\mathrm{A}} S^{\#}_{\mathrm{A}} \sigma_{\mathrm{AA}} \beta'_{\mathrm{A}}}{S^{\#}_{\mathrm{J}} + S^{\#}_{\mathrm{A}} + I^{\#}_{\mathrm{J}} + I^{\#}_{\mathrm{A}}} \cdot \dfrac{1}{\mu'_{\mathrm{A}}} \end{pmatrix}.
\end{aligned} \tag{A.13}$$

Elementary algebra of matrices gives the matrix-product form of $\mathbf{G}'$ in the main text.

If we denote the elements of $\mathbf{G}'$ by $a'_{ij}$ ($i, j$ run across A, J), its dominant eigenvalue (denoted $\Lambda[\mathbf{G}']$) is given by:

$$\Lambda[\mathbf{G}'] = \frac{a'_{\mathrm{JJ}} + a'_{\mathrm{AA}} + \sqrt{\left(a'_{\mathrm{JJ}} + a'_{\mathrm{AA}}\right)^2 - 4\left(a'_{\mathrm{JJ}} a'_{\mathrm{AA}} - a'_{\mathrm{JA}} a'_{\mathrm{AJ}}\right)}}{2}. \tag{A.14}$$

Here note that under weak selection (i.e., when $|\mathbf{v}' - \mathbf{v}|$ is negligibly small) and the continuity of $a'_{\mathrm{JJ}}$ and $a'_{\mathrm{AA}}$ with respect to $\mathbf{v}'$, we can show that:

$$a'_{\mathrm{JJ}} + a'_{\mathrm{AA}} < 2 \tag{A.15}$$

(see Appendix A.6; this inequality assures that the axis of symmetry of the characteristic function of $\mathbf{G}'$, which is a quadratic function, lies on the left of 1). With Eqn (A.14), we can consequently say that $\Lambda[\mathbf{G}'] > 1$ (the invadability condition) holds [1] if and only if:

$$w(\mathbf{v}', \mathbf{v}) := a'_{\mathrm{JJ}} + a'_{\mathrm{AA}} - \left(a'_{\mathrm{JJ}} a'_{\mathrm{AA}} - a'_{\mathrm{JA}} a'_{\mathrm{AJ}}\right) > 1. \tag{A.16}$$

Plugging Eqn (A.13) into Eqn (A.16) supplies:

$$w(\mathbf{v}', \mathbf{v}) = \alpha_{\mathrm{J}} \frac{S^{\#}_{\mathrm{J}}}{H^{\#}} \sigma_{\mathrm{JJ}} \frac{\beta'_{\mathrm{J}}}{\mu'_{\mathrm{J}}} + \frac{u}{\mu'_{\mathrm{J}}} \cdot \alpha_{\mathrm{J}} \frac{S^{\#}_{\mathrm{J}}}{H^{\#}} \sigma_{\mathrm{JA}} \frac{\beta'_{\mathrm{A}}}{\mu'_{\mathrm{A}}} + \alpha_{\mathrm{A}} \frac{S^{\#}_{\mathrm{A}}}{H^{\#}} \sigma_{\mathrm{AA}} \frac{\beta'_{\mathrm{A}}}{\mu'_{\mathrm{A}}} - \left(\sigma_{\mathrm{JJ}}\sigma_{\mathrm{AA}} - \sigma_{\mathrm{JA}}\sigma_{\mathrm{AJ}}\right) \frac{\alpha_{\mathrm{J}} S^{\#}_{\mathrm{J}} \alpha_{\mathrm{A}} S^{\#}_{\mathrm{A}}}{(H^{\#})^2} \cdot \frac{\beta'_{\mathrm{J}} \beta'_{\mathrm{A}}}{\mu'_{\mathrm{J}} \mu'_{\mathrm{A}}}. \tag{A.17}$$

Using the shorthand notation for $\pi'_{\mathrm{I}} = u/\mu'_{\mathrm{J}}$ (probability of successful maturation of juveniles infected

[1] The trick here is to isolate the square root on the left hand side and then square both sides.

by the mutant strain), $R'_X = \beta'_X/\mu'_X$ (the production from a X-stage host during its infectivity duration), $q^\#_{XY} := \alpha_X S^\#_X \sigma_{XY}/H^\#$ (the availability of stage-X hosts from the perspective of the parasite infecting a stage-Y hosts), and $\rho = \sigma_{JJ}\sigma_{AA} - \sigma_{JA}\sigma_{AJ}$ (assortativity), with all these substituted, one can recover the invasion fitness measure given in the main text (Eq 6).

An elementary calculation (using the endemic condition for the ODE, $\left(S^\#_J, S^\#_A, I^\#_J, I^\#_A\right)$) yields $w(\mathbf{v}, \mathbf{v}) \equiv 1$ for any $\mathbf{v}$; that is, the invasion fitness of a phenotypically neutral mutant is unity (and thus selectively neutral).

#### A.5 Selection gradient for adult virulence

Henceforth, by $f^\circ$, we mean that we evaluate a quantity $f$ at neutrality, $\mathbf{v}' = \mathbf{v}$. Partial differentiation of $w$ with respect to $v'_J, v'_A$ gives the selection gradient for the corresponding trait:

$$g_J(\mathbf{v}) = \left(\frac{\partial w(\mathbf{v}', \mathbf{v})}{\partial v'_J}\right)\bigg|_{\mathbf{v}'=\mathbf{v}}, \tag{A.18}$$

$$g_A(\mathbf{v}) = \left(\frac{\partial w(\mathbf{v}', \mathbf{v})}{\partial v'_A}\right)\bigg|_{\mathbf{v}'=\mathbf{v}}. \tag{A.19}$$

Upon some algebra, we get:

$$\begin{aligned} g_A(\mathbf{v}) = & \left\{\frac{\alpha_A S^\#_A \sigma_{AA}}{H^\#} \cdot \left(1 - \frac{\alpha_J S^\#_J \sigma_{JJ}}{H^\#} \cdot \frac{\beta_J}{\mu_J}\right) + \frac{\alpha_J S^\#_J \sigma_{JA}}{H^\#} \cdot \left(\frac{u}{\mu_J} + \frac{\alpha_A S^\#_A \sigma_{AJ}}{H^\#} \cdot \frac{\beta_J}{\mu_J}\right)\right\}^\circ \\ & \times \left(\frac{\beta_A}{\mu_A}\right)^\circ \cdot \left(\frac{1}{\beta_A} \cdot \frac{d\beta_A}{dv_A} - \frac{1}{\mu_A}\right)^\circ. \end{aligned} \tag{A.20}$$

It is only the final factor that can change its sign (see Footnote 2 in Appendix A.7). To obtain the selection gradient for juvenile virulence, more tedious work is needed. As such, we will use Fisher's reproductive value (Fisher 1958; Taylor 1990; Frank 1998; Caswell 2001).

#### A.6 Reproductive values

We here provide the reproductive-value based approach. Note that the case $\rho = 1$ violates this approach.

We shall first remember:

$$
\begin{aligned}
q^{\#}_{\mathrm{JJ}} &= \frac{\alpha_{\mathrm{J}} S^{\#}_{\mathrm{J}} \sigma_{\mathrm{JJ}}}{H^{\#}}, \\
q^{\#}_{\mathrm{JA}} &= \frac{\alpha_{\mathrm{J}} S^{\#}_{\mathrm{J}} \sigma_{\mathrm{JA}}}{H^{\#}}, \\
q^{\#}_{\mathrm{AJ}} &= \frac{\alpha_{\mathrm{A}} S^{\#}_{\mathrm{A}} \sigma_{\mathrm{AJ}}}{H^{\#}}, \\
q^{\#}_{\mathrm{AA}} &= \frac{\alpha_{\mathrm{A}} S^{\#}_{\mathrm{A}} \sigma_{\mathrm{AA}}}{H^{\#}}, \\
\pi'_{\mathrm{I}} &= \frac{u}{\mu'_{\mathrm{J}}}, \\
R'_{\mathrm{J}} &= \frac{\beta'_{\mathrm{J}}}{\mu'_{\mathrm{J}}}, \\
R'_{\mathrm{A}} &= \frac{\beta'_{\mathrm{A}}}{\mu'_{\mathrm{A}}};
\end{aligned}
\tag{A.21}
$$

then, we can get:

$$
\begin{aligned}
\mathbf{G}' &= \begin{pmatrix} a'_{\mathrm{JJ}} & a'_{\mathrm{JA}} \\ a'_{\mathrm{AJ}} & a'_{\mathrm{AA}} \end{pmatrix} \\
&= \begin{pmatrix} \frac{\alpha_{\mathrm{J}} S^{\#}_{\mathrm{J}} \sigma_{\mathrm{JJ}} \beta'_{\mathrm{J}}}{S^{\#}_{\mathrm{J}} + S^{\#}_{\mathrm{A}} + I^{\#}_{\mathrm{J}} + I^{\#}_{\mathrm{A}}} \cdot \frac{1}{\mu'_{\mathrm{J}}} + \frac{\alpha_{\mathrm{J}} S^{\#}_{\mathrm{J}} \sigma_{\mathrm{JA}} \beta'_{\mathrm{A}}}{S^{\#}_{\mathrm{J}} + S^{\#}_{\mathrm{A}} + I^{\#}_{\mathrm{J}} + I^{\#}_{\mathrm{A}}} \cdot \frac{u}{\mu'_{\mathrm{J}} \mu'_{\mathrm{A}}} & \frac{\alpha_{\mathrm{J}} S^{\#}_{\mathrm{J}} \sigma_{\mathrm{JA}} \beta'_{\mathrm{A}}}{S^{\#}_{\mathrm{J}} + S^{\#}_{\mathrm{A}} + I^{\#}_{\mathrm{J}} + I^{\#}_{\mathrm{A}}} \cdot \frac{1}{\mu'_{\mathrm{A}}} \\ \frac{\alpha_{\mathrm{A}} S^{\#}_{\mathrm{A}} \sigma_{\mathrm{AJ}} \beta'_{\mathrm{J}}}{S^{\#}_{\mathrm{J}} + S^{\#}_{\mathrm{A}} + I^{\#}_{\mathrm{J}} + I^{\#}_{\mathrm{A}}} \cdot \frac{1}{\mu'_{\mathrm{J}}} + \frac{\alpha_{\mathrm{A}} S^{\#}_{\mathrm{A}} \sigma_{\mathrm{AA}} \beta'_{\mathrm{A}}}{S^{\#}_{\mathrm{J}} + S^{\#}_{\mathrm{A}} + I^{\#}_{\mathrm{J}} + I^{\#}_{\mathrm{A}}} \cdot \frac{u}{\mu'_{\mathrm{J}} \mu'_{\mathrm{A}}} & \frac{\alpha_{\mathrm{A}} S^{\#}_{\mathrm{A}} \sigma_{\mathrm{AA}} \beta'_{\mathrm{A}}}{S^{\#}_{\mathrm{J}} + S^{\#}_{\mathrm{A}} + I^{\#}_{\mathrm{J}} + I^{\#}_{\mathrm{A}}} \cdot \frac{1}{\mu'_{\mathrm{A}}} \end{pmatrix} \\
&= \begin{pmatrix} q^{\#}_{\mathrm{JJ}} R'_{\mathrm{J}} + \pi'_{\mathrm{I}} q^{\#}_{\mathrm{JA}} R'_{\mathrm{A}} & q^{\#}_{\mathrm{JA}} R'_{\mathrm{J}} \\ q^{\#}_{\mathrm{AJ}} R'_{\mathrm{J}} + \pi'_{\mathrm{I}} q^{\#}_{\mathrm{AA}} R'_{\mathrm{A}} & q^{\#}_{\mathrm{AA}} R'_{\mathrm{A}} \end{pmatrix}.
\end{aligned}
\tag{A.22}
$$

At neutrality,

$$
\mathbf{G}^{\circ} = \begin{pmatrix} q^{\#}_{\mathrm{JJ}} R^{\circ}_{\mathrm{J}} + \pi_{\mathrm{I}} q^{\#}_{\mathrm{JA}} R^{\circ}_{\mathrm{A}} & q^{\#}_{\mathrm{JA}} R^{\circ}_{\mathrm{A}} \\ q^{\#}_{\mathrm{AJ}} R^{\circ}_{\mathrm{J}} + \pi_{\mathrm{I}} q^{\#}_{\mathrm{AA}} R^{\circ}_{\mathrm{A}} & q^{\#}_{\mathrm{AA}} R^{\circ}_{\mathrm{A}} \end{pmatrix}. \tag{A.23}
$$

Since the eigenvalue of $\mathbf{G}^{\circ}$ is unity, premultiplying the left eigenvector $(\ell^{\circ}_{\mathrm{J}}, \ell^{\circ}_{\mathrm{A}})$ must return $(\ell^{\circ}_{\mathrm{J}}, \ell^{\circ}_{\mathrm{A}})$:

$$
\left(\ell^{\circ}_{\mathrm{J}}, \ell^{\circ}_{\mathrm{A}}\right) \begin{pmatrix} q^{\#}_{\mathrm{JJ}} R^{\circ}_{\mathrm{J}} + \pi_{\mathrm{I}} q^{\#}_{\mathrm{JA}} R^{\circ}_{\mathrm{A}} & q^{\#}_{\mathrm{JA}} R^{\circ}_{\mathrm{A}} \\ q^{\#}_{\mathrm{AJ}} R^{\circ}_{\mathrm{J}} + \pi_{\mathrm{I}} q^{\#}_{\mathrm{AA}} R^{\circ}_{\mathrm{A}} & q^{\#}_{\mathrm{AA}} R^{\circ}_{\mathrm{A}} \end{pmatrix} = \left(\ell^{\circ}_{\mathrm{J}}, \ell^{\circ}_{\mathrm{A}}\right). \tag{A.24}
$$

Although it is possible to analytically solve $(\ell^{\circ}_{\mathrm{J}}, \ell^{\circ}_{\mathrm{A}})$, it does not lead to a transparent expression. Therefore, we instead derive the following (equivalent) relation:

$$\left(\ell_{\mathrm{J}}^{\circ},\ell_{\mathrm{A}}^{\circ}\right)(\mathbf{G}^{\circ}-\mathbf{I})=\left(\ell_{\mathrm{J}}^{\circ},\ell_{\mathrm{A}}^{\circ}\right)\begin{pmatrix} q_{\mathrm{JJ}}^{\#}R_{\mathrm{J}}^{\circ}+\pi_{\mathrm{I}}^{\circ}q_{\mathrm{JA}}^{\#}R_{\mathrm{A}}^{\circ}-1 & q_{\mathrm{JA}}^{\#}R_{\mathrm{A}}^{\circ} \\ q_{\mathrm{AJ}}^{\#}R_{\mathrm{J}}^{\circ}+\pi_{\mathrm{I}}^{\circ}q_{\mathrm{AA}}^{\#}R_{\mathrm{A}}^{\circ} & q_{\mathrm{AA}}^{\#}R_{\mathrm{A}}^{\circ}-1 \end{pmatrix}=(0,0) \tag{A.25}$$

(where $\mathbf{I}$ is the identity matrix), which explicitly (in elements) reads:

$$\ell_{\mathrm{J}}^{\circ}\underbrace{\left(1-q_{\mathrm{JJ}}^{\#}R_{\mathrm{J}}^{\circ}-\pi_{\mathrm{I}}^{\circ}q_{\mathrm{JA}}^{\#}R_{\mathrm{A}}^{\circ}\right)}_{=1-a_{\mathrm{JJ}}^{\circ}}=\ell_{\mathrm{A}}^{\circ}\left(q_{\mathrm{AJ}}^{\#}R_{\mathrm{J}}^{\circ}+\pi_{\mathrm{I}}^{\circ}q_{\mathrm{AA}}^{\#}R_{\mathrm{A}}^{\circ}\right), \tag{A.26}$$

$$\ell_{\mathrm{A}}^{\circ}\underbrace{\left(1-q_{\mathrm{AA}}^{\#}R_{\mathrm{A}}^{\circ}\right)}_{=1-a_{\mathrm{AA}}^{\circ}}=\ell_{\mathrm{J}}^{\circ}q_{\mathrm{JA}}^{\#}R_{\mathrm{A}}^{\circ}. \tag{A.27}$$

Using Eqns (A.26) and (A.27), we can now prove Eqn (A.15). Indeed, because the right-hand sides of Eqns (A.26) and (A.27) are both positive (by the Perron-Frobenius theorem), so are the left-hand sides of Eqns (A.26) and (A.27), implying that $1-a_{\mathrm{JJ}}^{\circ}>0$[2] and $1-a_{\mathrm{AA}}^{\circ}>0$ (remember the definition of $a$'s; see Eqn (A.13)). Under weak selection,

$$1-a'_{\mathrm{JJ}}+1-a'_{\mathrm{AA}}=\underbrace{1-a_{\mathrm{JJ}}^{\circ}}_{>0}+\underbrace{1-a_{\mathrm{AA}}^{\circ}}_{>0}+\underbrace{\left(a_{\mathrm{JJ}}^{\circ}-a'_{\mathrm{JJ}}\right)+\left(a_{\mathrm{AA}}^{\circ}-a'_{\mathrm{AA}}\right)}_{=O(|\mathbf{v}'-\mathbf{v}|)}; \tag{A.28}$$

where $O(|\mathbf{v}'-\mathbf{v}|)$ represents the Landau's big-$O$ (for $|\mathbf{v}'-\mathbf{v}|\to 0$) such that the latter two terms both tend towards zero as $|\mathbf{v}'-\mathbf{v}|\to 0$. Since $a'_{\mathrm{JJ}}$ and $a'_{\mathrm{AA}}$ are both continuous functions of $\mathbf{v}'$[3], under sufficiently weak mutation, we can assure the left-hand side of Eqn (A.28) be positive.

#### A.7 Selection gradient for juvenile virulence

In terms of $q$, $R$ and $\pi_{\mathrm{I}}$, the invasion fitness reads:

$$w(\mathbf{v}',\mathbf{v})=q_{\mathrm{JJ}}^{\#}R'_{\mathrm{J}}+\pi'_{\mathrm{I}}q_{\mathrm{JA}}^{\#}R'_{\mathrm{A}}+q_{\mathrm{AA}}^{\#}R'_{\mathrm{A}}-q_{\mathrm{JJ}}^{\#}q_{\mathrm{AA}}^{\#}R'_{\mathrm{A}}R'_{\mathrm{J}}+q_{\mathrm{JA}}^{\#}q_{\mathrm{AJ}}^{\#}R'_{\mathrm{A}}R'_{\mathrm{J}}. \tag{A.29}$$

Specifically, the fitness subcomponents involving $v'_{\mathrm{J}}$ amount to[4]:

$$\begin{aligned} w_{\mathrm{J}}(v'_{\mathrm{J}},v_{\mathrm{J}}) &:= q_{\mathrm{JJ}}^{\#}R'_{\mathrm{J}}+\pi'_{\mathrm{I}}q_{\mathrm{JA}}^{\#}R_{\mathrm{A}}^{\circ}-q_{\mathrm{JJ}}^{\#}q_{\mathrm{AA}}^{\#}R_{\mathrm{A}}^{\circ}R'_{\mathrm{J}}+q_{\mathrm{JA}}^{\#}q_{\mathrm{AJ}}^{\#}R_{\mathrm{A}}^{\circ}R'_{\mathrm{J}} \\ &= q_{\mathrm{JJ}}^{\#}\left(1-q_{\mathrm{AA}}^{\#}R_{\mathrm{A}}^{\circ}\right)R'_{\mathrm{J}}+\left(\pi'_{\mathrm{I}}+q_{\mathrm{AJ}}^{\#}R'_{\mathrm{J}}\right)q_{\mathrm{JA}}^{\#}R_{\mathrm{A}}^{\circ}, \end{aligned} \tag{A.30}$$

[2]This inequality consequently assures $1-\frac{\alpha_{\mathrm{J}}S_{\mathrm{J}}^{\#}\sigma_{\mathrm{JJ}}\beta_{\mathrm{J}}^{\circ}}{H^{\#}\mu_{\mathrm{J}}^{\circ}}=1-q_{\mathrm{JJ}}^{\#}R_{\mathrm{J}}^{\circ}>0$ when $\rho\neq 1$.

[3]Note here that $S_{\mathrm{J}}^{\#}, S_{\mathrm{A}}^{\#}, I_{\mathrm{J}}^{\#}$, and $I_{\mathrm{A}}^{\#}$ are all independent of $\mathbf{v}'$ because mutation is rare.

[4]Essencially, the invasion fitness subcomponents that do not contribute to the reproductive success of a parasite infecting a juvenile host are "excluded" from $w_{\mathrm{J}}$.

which we have evaluated at $v'_\mathrm{A} = v_\mathrm{A}$. Since $q^\#_\mathrm{JA} R^\circ_\mathrm{A} = \ell^\circ_\mathrm{A}/\ell^\circ_\mathrm{J}(1 - q^\#_\mathrm{AA} R^\circ_\mathrm{A})$, we finally have:

$$w_\mathrm{J}(v'_\mathrm{J}, v_\mathrm{J}) = \left(1 - q^\#_\mathrm{AA} R^\circ_\mathrm{A}\right)\left(q^\#_\mathrm{JJ} R'_\mathrm{J} + \frac{\ell^\circ_\mathrm{A}}{\ell^\circ_\mathrm{J}} \cdot \left(\pi'_\mathrm{I} + q^\#_\mathrm{AJ} R'_\mathrm{J}\right)\right) \tag{A.31}$$

when $\rho \neq 1$.

This expression is easier to differentiate:

$$\begin{aligned} g_\mathrm{J}(\mathbf{v}) &= \frac{1 - q^\#_\mathrm{AA} R^\circ_\mathrm{A}}{\ell^\circ_\mathrm{J}} \cdot \left(\ell^\circ_\mathrm{J} q^\#_\mathrm{JJ} R^\circ_\mathrm{J} \left(\frac{1}{\beta_\mathrm{J}} \cdot \frac{\mathrm{d}\beta_\mathrm{J}}{\mathrm{d}v_\mathrm{J}} - \frac{1}{\mu_\mathrm{J}}\right)^\circ + \ell^\circ_\mathrm{A} q^\#_\mathrm{AJ} R^\circ_\mathrm{J} \left(\frac{1}{\beta_\mathrm{J}} \cdot \frac{\mathrm{d}\beta_\mathrm{J}}{\mathrm{d}v_\mathrm{J}} - \frac{1}{\mu_\mathrm{J}}\right)^\circ - \ell^\circ_\mathrm{A} \pi^\circ_\mathrm{I} \cdot \frac{1}{\mu_\mathrm{J}}\right) \\ &= \frac{1 - q^\#_\mathrm{AA} R^\circ_\mathrm{A}}{\ell^\circ_\mathrm{J}} \left(\left(\ell^\circ_\mathrm{J} q^\#_\mathrm{JJ} + \ell^\circ_\mathrm{A} q^\#_\mathrm{AJ}\right) \times R^\circ_\mathrm{J} \left(\frac{1}{\beta_\mathrm{J}} \cdot \frac{\mathrm{d}\beta_\mathrm{J}}{\mathrm{d}v_\mathrm{J}} - \frac{1}{\mu_\mathrm{J}}\right)^\circ - \ell^\circ_\mathrm{A} \pi^\circ_\mathrm{I} \frac{1}{\mu_\mathrm{J}}\right). \end{aligned} \tag{A.32}$$

Using Eqn (A.27) to replace $\ell^\circ_\mathrm{A}$ (on the final factor) with $\left(\ell^\circ_\mathrm{J} q^\#_\mathrm{JA} + \ell^\circ_\mathrm{A} q^\#_\mathrm{AA}\right) R^\circ_\mathrm{A}$ we get the selection gradient of juvenile-virulence in the main text.

For the completeness, we can similarly get:

$$g_\mathrm{A}(\mathbf{v}) = \left(1 - q^\#_\mathrm{JJ} R^\circ_\mathrm{J}\right)\left(q^\#_\mathrm{AA} + \frac{\ell^\circ_\mathrm{J}}{\ell^\circ_\mathrm{A}} q^\#_\mathrm{JA}\right) \frac{\beta_\mathrm{A}}{\mu_\mathrm{A}} \left(\frac{1}{\beta_\mathrm{A}} \frac{\mathrm{d}\beta_\mathrm{A}}{\mathrm{d}v_\mathrm{A}} - \frac{1}{\mu_\mathrm{A}}\right). \tag{A.33}$$

Note that the first multiplicative term is always positive (see Footnote 2 and eqn (A.28)).

Using $\beta_\mathrm{A}(v_\mathrm{A}) = b_\mathrm{A} k_\mathrm{A} v_\mathrm{A}/(1 + k_\mathrm{A} v_\mathrm{A})$, it immediately follows that $g_\mathrm{A} = 0$ is solved by $v^*_\mathrm{A} = \sqrt{(m_\mathrm{A} + \gamma_\mathrm{A})/k_\mathrm{A}}$ and thus $v^*_\mathrm{A} = \sqrt{m_\mathrm{A}/k_\mathrm{A}}$ in the absence of recovery ($\gamma_\mathrm{A} = 0$).

##### A.8 Graph-theoretical approach

We here employ the graph-theoretical approach (GTA) developed by de Camino Beck & Lewis (2007), de Camino Beck & Lewis (2008), and de Camino Beck *et al.* (2008) to derive the invasion fitness measure (SI Fig 2), thereby checking the validity of $w(\mathbf{v}', \mathbf{v})$ in the main text. We write $\mathcal{R}'_\mathrm{m}$ for the invasion fitness derived through GTA.

The premise of the approach is to decompose fecundity output and state-transitions as in the next-generation theorem. The Jacobian around the endemic equilibrium reads:

$$\mathbf{J}' = \begin{pmatrix} q^\#_\mathrm{JJ}\beta'_\mathrm{J} - \mu'_\mathrm{J} & q^\#_\mathrm{JA}\beta'_\mathrm{A} \\ u + q^\#_\mathrm{AJ}\beta'_\mathrm{J} & q^\#_\mathrm{AA}\beta'_\mathrm{A} - \mu'_\mathrm{A} \end{pmatrix} = \begin{pmatrix} A'_\mathrm{JJ} & A'_\mathrm{JA} \\ A'_\mathrm{AJ} & A'_\mathrm{AA} \end{pmatrix} \tag{A.34}$$

de Camino Beck *et al.* (2008) defined three rules to algorithmically convert a compartmental structure into another; we detail these, partially borrowed from de Camino Beck *et al.* (2008), as follows:

**Rule A: Self-loop elimination (trivialization)** To reduce the loop $A'_\mathrm{XX}$ (which is $< 0$) to $-1$ at node X, every arc entering X has weight divided by $-A'_\mathrm{XX}\mathcal{R}'_\mathrm{m}$ (SI Fig 2A).

**Rule B: Parallel path elimination** For a path X → Y, if the path includes two weights then these are merged with the weight given as the sum of the two weights (SI Fig 2C).

**Rule C: trivial node elimination** For a trivialized node Y on a path X → Y → Z, the two arcs are replaced by a single arc X → Z with weight equal to $A'_{\mathrm{XY}}$ times $A'_{\mathrm{YZ}}$. Weights on multiple arcs X → Z are added. If there are no more paths through the trivial node Y, then it can be disregarded (SI Fig 2C).

Applying these rules, we can obtain the invasion condition, of:

$$\mathcal{R}'_{\mathrm{m}} = \sqrt{\frac{\pi'_{\mathrm{I}} + q^{\#}_{\mathrm{AJ}} R'_{\mathrm{J}}}{1 - q^{\#}_{\mathrm{JJ}} R'_{\mathrm{J}}} \cdot \frac{q^{\#}_{\mathrm{JA}} R'_{\mathrm{A}}}{1 - q^{\#}_{\mathrm{AA}} R'_{\mathrm{A}}}}, \tag{A.35}$$

with an elementary calculation showing $\mathcal{R}'_{\mathrm{m}} > 1$ if and only if $w(\mathbf{v}', \mathbf{v}) > 1$ under weak selection. The expression in Eqn (A.35) indicates that the invasion fitness can be decomposed into juvenile and adult components. Choice of $\mathcal{R}'_{\mathrm{m}}$, $w(\mathbf{v}', \mathbf{v})$, or $\Lambda[\mathbf{G}']$ is a matter of preference, all giving the same result for selection gradients and stability analyses. Taking advantage of deriving $\mathcal{R}'_{\mathrm{m}}$ (Eqn (A.35)), we can simplify the stability analysis (see the next subsection).

#### A.9 Attainability and Evolutionary stability

Here we outline the stability analyses for the evolutionary dynamics. Since the invasion fitness is not explicitly dependent on wild type strategy, the evolutionary stability and attainability conditions necessarily coincide (Otto & Day 2007). For this reason, we need only work on the evolutionary stability condition.

From Eqn (A.35), the invasion fitness is, in a product form, given by:

$$\mathcal{R}'_{\mathrm{m}}(\mathbf{v}') = \sqrt{\mathcal{W}'_{\mathrm{J}}\left(v'_{\mathrm{J}}\right) \cdot \mathcal{W}'_{\mathrm{A}}\left(v'_{\mathrm{A}}\right)}, \tag{A.36}$$

from which we can say that:

$$\begin{aligned} g_{\mathrm{J}} &\propto \left(\frac{\partial \mathcal{W}'_{\mathrm{J}}}{\partial v'_{\mathrm{J}}}\right)^{\circ}, \\ g_{\mathrm{A}} &\propto \left(\frac{\partial \mathcal{W}'_{\mathrm{A}}}{\partial v'_{\mathrm{A}}}\right)^{\circ}, \end{aligned} \tag{A.37}$$

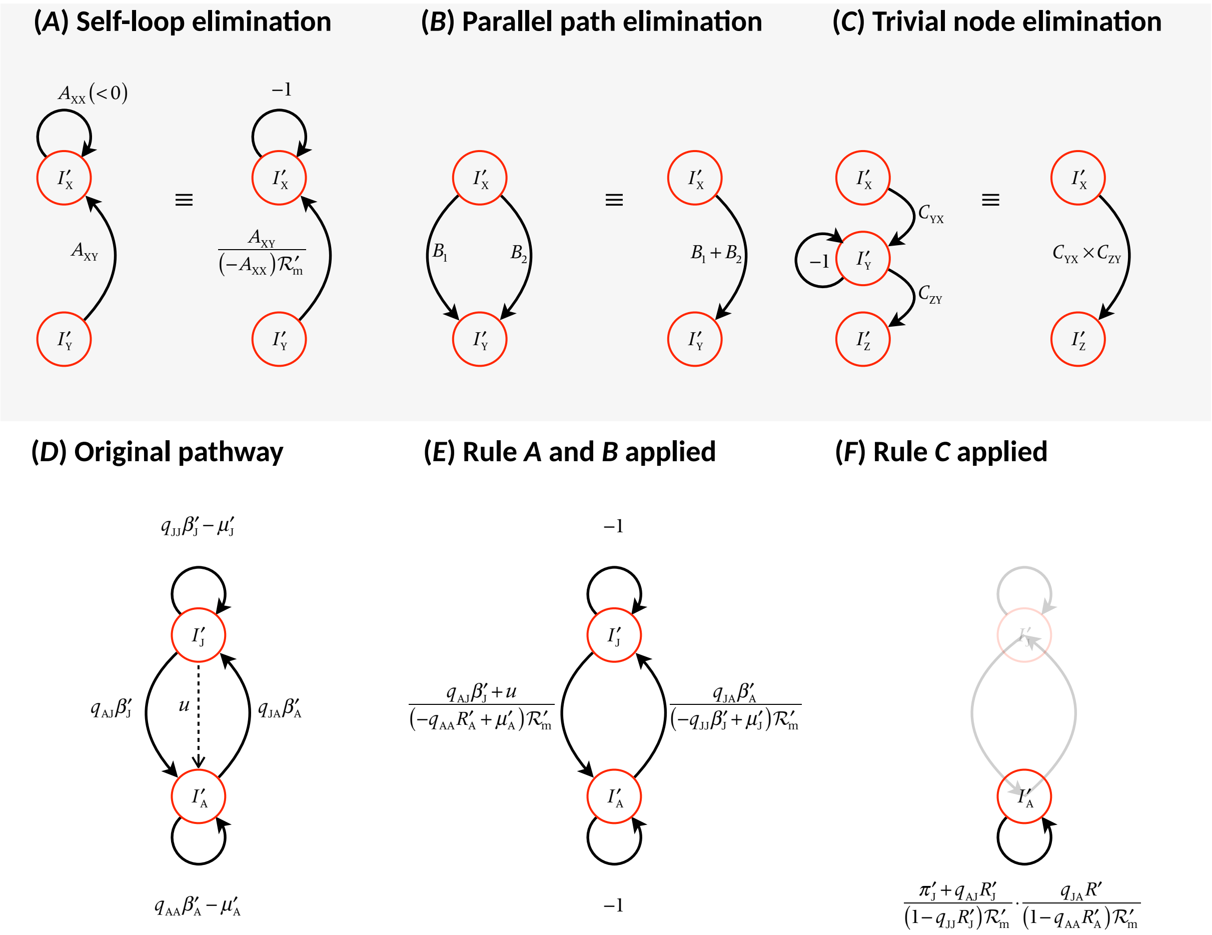

SI Figure 2: Graph-theoretical reduction of reproductive success pathways. $\mathcal{R}'_m$ represents the measure of invasion fitness; (A-C): general procedures in de Camino Beck & Lewis (2008). From this, by setting the last quantity unity, analytical expression of $\mathcal{R}'_m$ derives. (D): The "original" diagram depicting the pathways of reproductive success. $I'_J$ and $I'_A$ both have self-loop, so we will apply "self-loop" elimination rule. In addition, we apply parallel path elimination rule (by summing the transition, $u$, and the reproductive success of parasites infecting juveniles to adults through transmission, $W'_{AJ}$), obtaining (E): besides two trivial edges ("$-1$"), two nodes loop mutually and we apply node elimination rule, ending up with (F): the reproductive success of parasites infecting adults, the total number of "secondary" infection by mutant parasites, with all possible transmission-pathways included.

indicating that the Hessian matrix $\mathcal{H}$ of $\mathcal{R}'_{\mathrm{m}}$ at SS be given as a diagonal matrix; indeed:

$$\mathcal{H} = \begin{pmatrix} \dfrac{\partial^2 \mathcal{R}'_{\mathrm{m}}}{\partial {v'_{\mathrm{J}}}^2} & \dfrac{\partial^2 \mathcal{R}'_{\mathrm{m}}}{\partial v'_{\mathrm{J}} \partial v'_{\mathrm{A}}} \\ \dfrac{\partial^2 \mathcal{R}'_{\mathrm{m}}}{\partial v'_{\mathrm{A}} \partial v'_{\mathrm{J}}} & \dfrac{\partial^2 \mathcal{R}'_{\mathrm{m}}}{\partial {v'_{\mathrm{A}}}^2} \end{pmatrix} = \begin{pmatrix} \mathcal{W}'_{\mathrm{A}} \cdot \dfrac{\partial^2 \mathcal{W}'_{\mathrm{J}}}{\partial {v'_{\mathrm{J}}}^2} & \dfrac{\partial \mathcal{W}'_{\mathrm{J}}}{\partial v'_{\mathrm{J}}} \cdot \dfrac{\partial \mathcal{W}'_{\mathrm{A}}}{\partial v'_{\mathrm{A}}} \\ \dfrac{\partial \mathcal{W}'_{\mathrm{J}}}{\partial v'_{\mathrm{J}}} \cdot \dfrac{\partial \mathcal{W}'_{\mathrm{A}}}{\partial v'_{\mathrm{A}}} & \mathcal{W}'_{\mathrm{J}} \cdot \dfrac{\partial^2 \mathcal{W}'_{\mathrm{A}}}{\partial {v'_{\mathrm{A}}}^2} \end{pmatrix}, \tag{A.38}$$

which, evaluated at SS, gives a diagonal matrix because selection gradient vanishes at SS. Hence, it suffices to show that these diagonal terms – or the double partial derivatives – are both negative; that is, we shall show:

$$\begin{aligned} \left( \frac{\partial^2 \mathcal{W}'_{\mathrm{J}}}{\partial {v'_{\mathrm{J}}}^2} \right)^{\circ} &< 0, \\ \left( \frac{\partial^2 \mathcal{W}'_{\mathrm{A}}}{\partial {v'_{\mathrm{A}}}^2} \right)^{\circ} &< 0. \end{aligned} \tag{A.39}$$

[∵] First, Eqn (A.36) indicates:

$$\begin{aligned} \mathcal{W}'_{\mathrm{J}} &= \frac{\pi'_{\mathrm{I}} + q^{\#}_{\mathrm{AJ}} R'_{\mathrm{J}}}{1 - q^{\#}_{\mathrm{JJ}} R'_{\mathrm{J}}}, \\ \mathcal{W}'_{\mathrm{A}} &= \frac{q^{\#}_{\mathrm{JA}} R'_{\mathrm{A}}}{1 - q^{\#}_{\mathrm{AA}} R'_{\mathrm{A}}}, \end{aligned} \tag{A.40}$$

which with straightforward calculations gives:

$$\left. \frac{\partial^2 \mathcal{W}'_{\mathrm{A}}}{\partial {v'_{\mathrm{A}}}^2} \right|_{\mathbf{v}=\mathbf{v}^*} = q^*_{\mathrm{JA}} \times \left( \underbrace{\frac{\partial^2 R'_{\mathrm{A}}}{\partial {v'_{\mathrm{A}}}^2}}_{<0} \times \left(1 - q^*_{\mathrm{AA}} R^*_{\mathrm{A}}\right)^2 + 2 \underbrace{\left( \frac{\partial R'_{\mathrm{A}}}{\partial v'_{\mathrm{A}}} \right)^2}_{=0 \text{ at SS}} \left(1 - q^*_{\mathrm{AA}} R^*_{\mathrm{A}}\right) q^*_{\mathrm{AA}} \right) < 0, \tag{A.41}$$

as desired; note that if $\rho = 1$ then this second derivative is always null at the SS, meaning that any mutants in $v_{\mathrm{A}}$ are selectively neutral at the SS.

Second, the first derivative of $\mathcal{W}_{\mathrm{J}}'$ (prior to being evaluated at SS) reads:

$$\begin{aligned}\frac{\partial \mathcal{W}_{\mathrm{J}}'}{\partial v_{\mathrm{J}}'} &= \frac{1}{\left(1 - q_{\mathrm{JJ}}^{\#} R_{\mathrm{J}}'\right)^2} \left\{ \left(\pi_{\mathrm{I}}^{[1]} + q_{\mathrm{AJ}}^{\#} R_{\mathrm{J}}^{[1]}\right) \left(1 - q_{\mathrm{JJ}}^{\#} R_{\mathrm{J}}'\right) + q_{\mathrm{JJ}}^{\#} R_{\mathrm{J}}^{[1]} \left(\pi_{\mathrm{I}}' + q_{\mathrm{AJ}}^{\#} R_{\mathrm{J}}'\right) \right\} \\ &= \left\{ \left(\pi_{\mathrm{I}}^{[1]} + q_{\mathrm{AJ}}^{\#} R_{\mathrm{J}}^{[1]}\right) \left(1 - q_{\mathrm{JJ}}^{\#} R_{\mathrm{J}}'\right) + q_{\mathrm{JJ}}^{\#} R_{\mathrm{J}}^{[1]} \left(\pi_{\mathrm{I}}' + q_{\mathrm{AJ}}^{\#} R_{\mathrm{J}}'\right) \right\} \left(1 - q_{\mathrm{JJ}}^{\#} R_{\mathrm{J}}'\right)^{-2}\end{aligned} \tag{A.42}$$

(with the shorthand notation [1] for its first derivative with respect to $v_{\mathrm{J}}'$), from which, as the selection gradient $g_{\mathrm{J}}(\mathbf{v})$ vanishes at $\mathbf{v}^*$, we have:

$$\left(\pi_{\mathrm{I}}^{\circ} q_{\mathrm{JJ}}^{\#} + q_{\mathrm{AJ}}^{\#}\right) \left(R_{\mathrm{J}}^{[1]}\right)^{\circ} = -\left(\pi_{\mathrm{I}}^{[1]}\right)^{\circ} \left(1 - q_{\mathrm{JJ}}^{\#} R_{\mathrm{J}}'\right). \tag{A.43}$$

Also, using $R_{\mathrm{J}}' = \beta_{\mathrm{J}}'/\mu_{\mathrm{J}}'$, we immediately have:

$$\begin{aligned}\left(R_{\mathrm{J}}^{[1]}\right)^{\circ} &= \left(\frac{\mathrm{d}R_{\mathrm{J}}'}{\mathrm{d}v_{\mathrm{J}}'}\right)^{\circ} = \left(\frac{\beta_{\mathrm{J}}^{[1]} \mu_{\mathrm{J}} - \beta_{\mathrm{J}}}{\mu_{\mathrm{J}}^2}\right)^{\circ}, \\ \left(R_{\mathrm{J}}^{[2]}\right)^{\circ} &= \left(\frac{\mathrm{d}^2 R_{\mathrm{J}}'}{\mathrm{d}v_{\mathrm{J}}'^2}\right)^{\circ} = \left(\frac{\beta_{\mathrm{J}}^{[2]}}{\mu_{\mathrm{J}}} - \frac{2}{\mu_{\mathrm{J}}} R_{\mathrm{J}}^{[1]}\right)^{\circ}.\end{aligned} \tag{A.44}$$

The second derivative of $\mathcal{W}_{\mathrm{J}}'$ evaluated at SS reads:

$$\begin{aligned}\left.\frac{\partial^2 \mathcal{W}_{\mathrm{J}}'}{\partial v_{\mathrm{J}}'^2}\right|_{\mathbf{v}=\mathbf{v}^*} &= \left\{ \pi_{\mathrm{I}}^{[2]} \left(1 - q_{\mathrm{JJ}}^* R_{\mathrm{J}}^*\right) + \pi_{\mathrm{I}}^{[1]} \left(-q_{\mathrm{JJ}}^* R_{\mathrm{J}}^{[1]}\right) + q_{\mathrm{AJ}}^* R_{\mathrm{J}}^{[2]} + q_{\mathrm{JJ}}^* R_{\mathrm{J}}^{[2]} \pi_{\mathrm{I}} + q_{\mathrm{JJ}}^* R_{\mathrm{J}}^{[1]} \pi_{\mathrm{I}}^{[1]} \right\}^* \cdot \left(1 - q_{\mathrm{JJ}}^* R_{\mathrm{A}}^*\right)^{-2} \\ &\quad + 2 q_{\mathrm{JJ}}^* \left(R_{\mathrm{J}}^{[1]}\right)^* \cdot \left(1 - q_{\mathrm{JJ}}^* R_{\mathrm{J}}^*\right)^{-3} \cdot \underbrace{\left\{ \left(\pi_{\mathrm{I}}^{[1]} + q_{\mathrm{AJ}}^* R_{\mathrm{J}}^{[1]}\right) \left(1 - q_{\mathrm{JJ}}^* R_{\mathrm{J}}^*\right) + q_{\mathrm{JJ}}^* R_{\mathrm{J}}^{[1]} \left(\pi_{\mathrm{I}}^* + q_{\mathrm{AJ}}^* R_{\mathrm{J}}^*\right) \right\}^*}_{\propto g_{\mathrm{J}}(\mathbf{v}^*)=0 \text{ at SS}} \\ &= \left\{ \pi_{\mathrm{I}}^{[2]} \left(1 - q_{\mathrm{JJ}}^* R_{\mathrm{J}}^*\right) + q_{\mathrm{AJ}}^* R_{\mathrm{J}}^{[2]} + q_{\mathrm{JJ}}^* R_{\mathrm{J}}^{[2]} \pi_{\mathrm{I}} \right\}^* \cdot \left(1 - q_{\mathrm{JJ}}^* R_{\mathrm{J}}^*\right)^{-2}\end{aligned} \tag{A.45}$$

As $\pi_{\mathrm{I}} = u/(u + m_{\mathrm{J}} + v_{\mathrm{J}})$, the first and second derivatives at SS are given by:

$$\left(\pi_{\mathrm{I}}^{[1]}\right)^{\circ} = -\left(\frac{\pi_{\mathrm{I}}}{\mu_{\mathrm{J}}}\right)^{\circ}, \tag{A.46}$$

$$\left(\pi_{\mathrm{I}}^{[2]}\right)^{\circ} = \left(\frac{2\pi_{\mathrm{I}}}{(\mu_{\mathrm{J}})^2}\right)^{\circ}, \tag{A.47}$$

which, plugged into Eqn (A.45), give:

$$
\begin{aligned}
\left.\frac{\partial^2 \mathcal{W}_{\mathrm{J}}'}{\partial v_{\mathrm{J}}'^2}\right|_{\mathbf{v}^*} &= \left\{\pi_{\mathrm{I}}^{[2]}\left(1-q_{\mathrm{JJ}}^{\#}R_{\mathrm{J}}\right)+q_{\mathrm{AJ}}^{\#}R_{\mathrm{J}}^{[2]}+q_{\mathrm{JJ}}^{\#}R_{\mathrm{J}}^{[2]}\pi_{\mathrm{I}}\right\}^{\circ}\cdot\left(1-q_{\mathrm{JJ}}^{\#}R_{\mathrm{J}}^{\circ}\right)^{-2} \\
&= \left\{\pi_{\mathrm{I}}^{[2]}\left(1-q_{\mathrm{JJ}}^{\#}R_{\mathrm{J}}\right)+\left(\pi_{\mathrm{I}}q_{\mathrm{JJ}}^{\#}+q_{\mathrm{AJ}}^{\#}\right)\underbrace{R_{\mathrm{J}}^{[2]}}_{\text{use Eqn (A.44)}}\right\}^{\circ}\cdot\left(1-q_{\mathrm{JJ}}^{\#}R_{\mathrm{J}}^{\circ}\right)^{-2} \\
&= \left\{\underbrace{\pi_{\mathrm{I}}^{[2]}}_{\text{use Eqn (A.47)}}\left(1-q_{\mathrm{JJ}}^{\#}R_{\mathrm{J}}\right)+\left(\pi_{\mathrm{I}}q_{\mathrm{JJ}}^{\#}+q_{\mathrm{AJ}}^{\#}\right)\left(\frac{\beta_{\mathrm{J}}^{[2]}}{\mu_{\mathrm{J}}}-\frac{2}{\mu_{\mathrm{J}}}R_{\mathrm{J}}^{[1]}\right)\right\}^{\circ}\cdot\left(1-q_{\mathrm{JJ}}^{\#}R_{\mathrm{J}}^{\circ}\right)^{-2} \\
&= \left\{\frac{2\pi_{\mathrm{I}}}{\left(\mu_{\mathrm{J}}\right)^2}\left(1-q_{\mathrm{JJ}}^{\#}R_{\mathrm{J}}\right)+\left(\pi_{\mathrm{I}}q_{\mathrm{JJ}}^{\#}+q_{\mathrm{AJ}}^{\#}\right)\left(\frac{\beta_{\mathrm{J}}^{[2]}}{\mu_{\mathrm{J}}}-\underbrace{\frac{2}{\mu_{\mathrm{J}}}R_{\mathrm{J}}^{[1]}}_{\text{use Eqn (A.44)}}\right)\right\}^{\circ}\cdot\left(1-q_{\mathrm{JJ}}^{\#}R_{\mathrm{J}}^{\circ}\right)^{-2} \\
&= \left\{\frac{2\pi_{\mathrm{I}}}{\left(\mu_{\mathrm{J}}\right)^2}\left(1-q_{\mathrm{JJ}}^{\#}R_{\mathrm{J}}\right)+\left(\pi_{\mathrm{I}}q_{\mathrm{JJ}}^{\#}+q_{\mathrm{AJ}}^{\#}\right)\left(\frac{\beta_{\mathrm{J}}^{[2]}}{\mu_{\mathrm{J}}}\right)+\underbrace{\frac{2\pi_{\mathrm{I}}^{[1]}}{\mu_{\mathrm{J}}}}_{=-2\pi_{\mathrm{I}}/\mu_{\mathrm{J}}^2}\left(1-q_{\mathrm{JJ}}^{\#}R_{\mathrm{J}}\right)\right\}^{\circ}\cdot\left(1-q_{\mathrm{JJ}}^{\#}R_{\mathrm{J}}^{\circ}\right)^{-2} \\
&= \underbrace{\left(\pi_{\mathrm{I}}^{\circ}q_{\mathrm{JJ}}^{\#}+q_{\mathrm{AJ}}^{\#}\right)}_{>0}\underbrace{\left(\frac{\beta_{\mathrm{J}}^{[2]}}{\mu_{\mathrm{J}}}\right)^{\circ}}_{<0}\cdot\left(1-q_{\mathrm{JJ}}^{\#}R_{\mathrm{J}}^{\circ}\right)^{-2}<0,
\end{aligned}
\tag{A.48}
$$

which completes the proof of the statement Eqn (A.39).

#### A.10 Condition for parasite persistence

In the absence of diseases,

$$
\begin{aligned}
\frac{\mathrm{d}S_{\mathrm{J}}}{\mathrm{d}t} &= \left(r-\kappa S_{\mathrm{A}}\right)\cdot S_{\mathrm{A}}-\left(u+m_{\mathrm{J}}\right)S_{\mathrm{J}}, \\
\frac{\mathrm{d}S_{\mathrm{A}}}{\mathrm{d}t} &= uS_{\mathrm{J}}-m_{\mathrm{A}}S_{\mathrm{A}}.
\end{aligned}
\tag{A.49}
$$

Disease-free equilibrium is given by:

$$\left(S_{\mathrm{J}}, S_{\mathrm{A}}\right) = \left(S_{\mathrm{J}}^{(0)}, S_{\mathrm{A}}^{(0)}\right) = \left(\frac{m_{\mathrm{A}}}{u} \cdot \frac{r - m_{\mathrm{A}}\dfrac{u + m_{\mathrm{J}}}{u}}{\kappa}, \frac{r - m_{\mathrm{A}}\dfrac{u + m_{\mathrm{J}}}{u}}{\kappa}\right), \tag{A.50}$$

from which we can get:

$$\frac{S_{\mathrm{A}}^{(0)}}{S_{\mathrm{J}}^{(0)} + S_{\mathrm{A}}^{(0)}} = \frac{u}{u + m_{\mathrm{A}}}. \tag{A.51}$$

Parasites attempting to invade such a disease-free, stage-structured host population can establish only if:

$$R_0 = \alpha_{\mathrm{J}} \frac{m_{\mathrm{A}}}{u + m_{\mathrm{A}}} \sigma_{\mathrm{JJ}} \frac{\beta_{\mathrm{J}}}{\mu_{\mathrm{J}}} + \frac{u}{\mu_{\mathrm{J}}} \cdot \alpha_{\mathrm{J}} \frac{m_{\mathrm{A}}}{u + m_{\mathrm{A}}} \sigma_{\mathrm{JA}} \frac{\beta_{\mathrm{A}}}{\mu_{\mathrm{A}}} + \alpha_{\mathrm{A}} \frac{u}{u + m_{\mathrm{A}}} \sigma_{\mathrm{AA}} \frac{\beta_{\mathrm{A}}}{\mu_{\mathrm{A}}} - \alpha_{\mathrm{J}} \alpha_{\mathrm{A}} \rho \cdot \frac{u}{u + m_{\mathrm{A}}} \cdot \frac{m_{\mathrm{J}}}{u + m_{\mathrm{A}}} \cdot \frac{\beta_{\mathrm{J}} \beta_{\mathrm{A}}}{\mu_{\mathrm{J}} \mu_{\mathrm{A}}} > 1. \tag{A.52}$$

When the outcomes of selection (i.e., $(v_{\mathrm{J}}, v_{\mathrm{A}}) = (v_{\mathrm{J}}^*, v_{\mathrm{A}}^*)$) violate this condition, parasite extinction (evolutionary suicide) can occur.

##### A.11 When $\rho = 1$ (fully assortative transmission)

Finally, we detail what if $\rho = 1$; then $\Lambda\left[\mathbf{G}'\right]$ is given by:

$$\Lambda\left[\mathbf{G}'\right] = \max\left(q_{\mathrm{JJ}}^{\#} R_{\mathrm{J}}', q_{\mathrm{AA}}^{\#} R_{\mathrm{A}}'\right) = \max\left(\frac{R_{\mathrm{J}}'}{R_{\mathrm{J}}}, \frac{R_{\mathrm{A}}'}{R_{\mathrm{A}}}\right) \tag{A.53}$$

In this case, obtaining the selection gradient is not needed. Instead, we can directly see that the evolutionary stability condition reads:

$$\max\left(q_{\mathrm{JJ}}^{\#} R_{\mathrm{J}}', q_{\mathrm{AA}}^{\#} R_{\mathrm{A}}'\right) = \max\left(\frac{R_{\mathrm{J}}'}{R_{\mathrm{J}}}, \frac{R_{\mathrm{A}}'}{R_{\mathrm{A}}}\right) < 1 \tag{A.54}$$

for any $\mathbf{v}' \neq \mathbf{v}$. This is thus obtained by jointly maximizing two functions $R_{\mathrm{J}}' = \beta_{\mathrm{J}}(v_{\mathrm{J}}')/\mu_{\mathrm{J}}'$ and $R_{\mathrm{A}}' = \beta_{\mathrm{A}}(v_{\mathrm{A}}')/\mu_{\mathrm{A}}'$, giving the CSS as $\left(v_{\mathrm{J}}^*, v_{\mathrm{A}}^*\right) = \left(\sqrt{(m_{\mathrm{J}} + u)/k_{\mathrm{J}}}, \sqrt{(m_{\mathrm{A}}/k_{\mathrm{A}})}\right)$.

#### B Robustness

In the main text, we have assumed:

- There is no recovery: $\gamma_{\mathrm{J}} = \gamma_{\mathrm{A}} = 0$;
- Susceptibility is the same: $\alpha_{\mathrm{J}} = \alpha_{\mathrm{A}} = 1$;
- Maximum infectiousness is the same: $b_{\mathrm{J}} = b_{\mathrm{J}} = 10$;

- The response of infectiousness to increased virulence (i.e., the efficiency improved growth due to exploitation) is the same: $k_\mathrm{J} = k_\mathrm{A} = 1$;
- Transmission is frequency-dependent: $\phi_\mathrm{XY} = \alpha_\mathrm{X}\sigma_\mathrm{XY}\beta_\mathrm{Y}I_\mathrm{Y}/H^\#$.
- Fecundity is the same for susceptible and infected adults.

Here we will check the robustness of our prediction against these variants. Specifically, we will work on the specificity in:

- recovery: $(\gamma_\mathrm{J}, \gamma_\mathrm{A})$;
- susceptibility: $(\alpha_\mathrm{J}, \alpha_\mathrm{A})$;
- tolerance: $(k_\mathrm{J}, k_\mathrm{A})$;
- resistance: $(b_\mathrm{J}, b_\mathrm{A})$;
- density-dependent transmission: $\phi_\mathrm{XY} = \alpha_\mathrm{X}\sigma_\mathrm{XY}\beta_\mathrm{Y}I_\mathrm{Y}$.
- fecundity changes in infected adults, $1 - h$ (with $h$ possibly negative).

Note that we did not always show the full range of $\rho \in [-1, 1]$ and $\theta_\mathrm{A} \in [0, 1]$, because the numerical routines are computationally expensive. Also, we used the default parameter values unless otherwise specified; specifically, $m_\mathrm{J} = m_\mathrm{A} = 1$.

##### B.1 Recovery

We used relatively small values of $(\gamma_\mathrm{J}, \gamma_\mathrm{A})$ in the ODE, because high recovery can readily result in parasite extinction. We again numerically obtained the CSS virulence and plotted them on the $(\rho, u)$-plane. We can see that our prediction is qualitatively robust against this variant. Quantitative differences are that recovery can in general favour fast exploitation, which is obvious from the CSS for adult virulence, $v_\mathrm{A}^* = \sqrt{(m_\mathrm{A} + \gamma_\mathrm{A})/k_\mathrm{A}} > \sqrt{m_\mathrm{A}/k_\mathrm{A}}$. In the numerical examble, $\gamma_\mathrm{A} = 0.25, k_\mathrm{A} = m_\mathrm{A} = 1$ yields $v_\mathrm{A}^* = \sqrt{5}/2 \approx 1.118$. As for juvenile virulence $v_\mathrm{J}^*$, the general trend is unchanged (SI Fig 5).

As recovery increases, evolutionary suicide is more readily to occur (white zone). This is so because parasites have to faster exploit the hosts while there is no trade-off between recovery and other traits (i.e., other traits do not compensate the decreased infectious period).

Overall, the effects of recovery are similar to those of mortality (see Figure 2 in the main text).

##### B.2 Susceptibility

We here introduce a difference in $\alpha$'s, which corresponds to the situation where juveniles and adults show quantitatively different transmission-blocking mechanisms. This does not affect the results critically; a difference is that evolutionary suicide is more likely to occur with smaller $\alpha$'s.

##### B.3 Tolerance

Tolerance, or reduced negative impacts of the disease on hosts, can affect the tradeoff through $k_\mathrm{X}$. For simplicity, we assume that $b_\mathrm{X}$ is constant (see next section). To incorporate tolerance, we further decompose parasite-induced mortality into $v_\mathrm{X} = (1 - \tau_\mathrm{X})\, e_\mathrm{X}$, where $\tau_\mathrm{X}$ tunes tolerance and $e_\mathrm{X}$ represents exploitation. Infectiousness-exploitation tradeoff can be given by:

$$\begin{aligned}\beta_X(e_X) &= b_X \frac{k_X e_X}{1 + k_X e_X} \\ &= b_X \frac{\frac{k_X}{1-\tau_X} v_X}{1 + \frac{k_X}{1-\tau_X} v_X},\end{aligned} \tag{B.55}$$

whereas a derivative is given by:

$$\frac{\mathrm{d}v_X}{\mathrm{d}e_X} = 1 - \tau_X, \tag{B.56}$$

which is a constant for each X (with X = J or A). Higher tolerance (larger $\tau_X$) leads to larger $k_X/(1 - \tau_X)$.

Marginal value theorem (Charnov 1976) shows that SS solves:

$$\frac{1}{\beta_A} \cdot \frac{\mathrm{d}\beta_A}{\mathrm{d}e_A} = \frac{1 - \tau_A}{(1 - \tau_A)e_A + m_A}, \tag{B.57}$$

supplying $e_A^* = \sqrt{m_A(1 - \tau_A)/k_A}$. Hence SS for $e_A$ is smaller with tolerance. To look at the consequences for $e_J$, we again solved the equations, observing that the results are qualitatively unchanged.

##### B.4 Infectiousness

We assess the effects of varying $b_X$. Obviously, increasing $b_X$ results in higher transmission but does not affect the SS for adult virulence (SI Fig 6).

##### B.5 Density-dependent transmission

Because the densities would be of greater importance to the force of infection with this assumption, we used a smaller value of $b_J = b_A = 0.13$. We found quantitatively similar outcomes (SI Fig 7).

##### B.6 Fecundity virulence and evolutionarily stable resource shifts

We here explore the effects of fecundity shifts on evolution of virulence, looking at the possibility that parasites deprive some amounts of resource of infected hosts that would have been otherwise available to the hosts for reproduction. We do so by considering two models: in the first model, we assume that the fecundity shift in adults, denoted $h$, is a constant ($h$ can be negative). We consequently found that the results are robust.

#### C Generalized pathway structure

In the main text we posed three constraints, namely normalization ($\sigma_{JA} + \sigma_{AA} = 1$ and $\sigma_{AJ} + \sigma_{JJ} = 1$) and symmetry ($\sigma_{AJ} = \sigma_{JA}$), thereby tuning a single parameter of the diagonal element ($\sigma_{JJ} = \sigma_{AA} = \sigma$ was the parameter of interest). Here we relax each of these assumptions, which we found did not dramatically change our predictions.

We first of all remark that transmission terms are governed by four compound quantities $\phi_{\mathrm{XY}}$ (with X and Y running across J and A), meaning that eight (or two pairs of four) multiplicative terms for $\alpha_{\mathrm{J}}, \alpha_{\mathrm{A}}, b_{\mathrm{J}}, b_{\mathrm{A}}, \sigma_{\mathrm{JJ}}, \sigma_{\mathrm{AJ}}, \sigma_{\mathrm{JA}}, \sigma_{\mathrm{AA}}$ are redundant; we can impose four constraints to these parameters. For instance, the condition $\sigma_{\mathrm{AA}} = 1 \gg \sigma_{\mathrm{JJ}} = 1/100$ with $\sigma_{\mathrm{JA}} = \sigma_{\mathrm{AJ}} = 1/2$ with $\alpha_{\mathrm{J}} = \alpha_{\mathrm{A}} = 1$ and $b_{\mathrm{J}} = b_{\mathrm{A}} = 10$ (such that pathway is symmetric), is equivalent to $\sigma_{\mathrm{AA}} = 2/3, \sigma_{\mathrm{JA}} = 1/3, b_{\mathrm{A}} = 15, \sigma_{\mathrm{AJ}} = 50/51, \sigma_{\mathrm{JJ}} = 1/51, b_{\mathrm{J}} = 51/10$ (such that, with $\sigma_{\mathrm{AJ}} + \sigma_{\mathrm{JJ}} = \sigma_{\mathrm{AA}} + \sigma_{\mathrm{JA}} = 1$, the pathway pattern is normalized). The product-decomposition of the force of infection $\phi_{\mathrm{XY}}$ is thus not unique. Therefore, as we have already shown that neither does a slight difference in susceptibility $\alpha_{\mathrm{J}} \neq \alpha_{\mathrm{A}}$ or in infectiousness $b_{\mathrm{A}} \neq b_{\mathrm{A}}$ affect the results, we can restrict ourselves to $\alpha_{\mathrm{J}} = \alpha_{\mathrm{A}}$ and $b_{\mathrm{J}} = b_{\mathrm{A}}$ (with two parameters reduced).

We further impose more constraints. In the first case, we assume $\sigma_{\mathrm{JJ}} = 1 - \sigma_{\mathrm{AJ}}$ and $\sigma_{\mathrm{AA}} = 1 - \sigma_{\mathrm{JA}}$ (normalized pathway) and varying $\sigma_{\mathrm{JJ}}$ and $\sigma_{\mathrm{AA}}$; the second is to fix $\sigma_{\mathrm{AA}}$ and vary $\sigma_{\mathrm{JJ}}$ and $\sigma_{\mathrm{JA}} = \sigma_{\mathrm{AJ}}$ (symmetric pathway).

###### "Normalized pathway" : varying $\sigma_{\mathrm{JJ}} = 1 - \sigma_{\mathrm{AJ}}$ and $\sigma_{\mathrm{AA}} = 1 - \sigma_{\mathrm{JA}}$

The pathway matrix reads

$$\begin{pmatrix} \sigma_{\mathrm{JJ}} & 1-\sigma_{\mathrm{JJ}} \\ 1-\sigma_{\mathrm{AA}} & \sigma_{\mathrm{AA}} \end{pmatrix}, \tag{C.58}$$

with $\rho = \sigma_{\mathrm{JJ}} + \sigma_{\mathrm{AA}} - 1 \in [-1, 1]$. SI Fig 9 suggests that when $\theta_{\mathrm{A}}$ is small, higher assortativity (top right zone) favors higher juveniles virulence (top panels) but this trend turns over as $\theta_{\mathrm{A}}$ becomes larger.

###### "Symmetric pathway" : varying $\sigma_{\mathrm{JJ}}$ and $\sigma_{\mathrm{AJ}} = \sigma_{\mathrm{JA}}$, with $\sigma_{\mathrm{AA}}$ fixed

The pathway matrix reads

$$\begin{pmatrix} \sigma_{\mathrm{JJ}} & \sigma_{\mathrm{AJ}} \\ \sigma_{\mathrm{AJ}} & \sigma_{\mathrm{AA}} \end{pmatrix}, \tag{C.59}$$

with $\rho = \sigma_{\mathrm{JJ}} \cdot \sigma_{\mathrm{AA}} - \sigma_{\mathrm{AJ}}^2 \in [-1, 1]$. SI Fig 10 shows that, when $\theta_{\mathrm{A}}$ is small (or large), smaller (or larger) $\sigma_{\mathrm{AJ}}$ favors higher juvenile-virulence (respectively). Therefore, the transmission pathway interpretation is again consistent and thus robust to this variant.

### D Empirical data figure and credits

We conducted several literature searches in Google Scholar combining the terms "age- related" or "age-dependent" or "stage-dependent" or "juvenile" + "susceptibility" or "resistance" or "tolerance" or "immunocompetence" + "infection" or "infectious disease". From these searches, we collected data from papers where the parasite could be judged to be adapted to its host (ie. not a recent host shift and without significant multi- species transmission) and where differences in virulence across life stages could be distinguished from age-related trends in additional mortality due to increasing adaptive immunity with age due to previous exposure and increased mortality of poor-condition hosts during the juvenile stages. Therefore, we collected data from papers for host- pathogen systems where adaptive immunity to the pathogen was not significant or infection-related mortality was measured in naïve juveniles and adults in either a natural population or in an experimental lab population. From the papers that we found, we also searched their citations and papers that cited them for other publications that we may have missed in the first search. After we had found papers with reliable data on age- biased virulence, we searched for "host" and "life history" or "age at reproduction" to find data on the host's maturation rate. Finally, we searched for transmission assortativity data for each selected system by searching the terms "host"+ "transmission" or "contact network"+ "age" or "stage" or "juvenile". We used estimated values of $v_{\mathrm{J}}$ versus $v_{\mathrm{A}}$. The extracted data are plotted against a $(\rho, \theta_{\mathrm{A}})$-plane.

Concerning the data on asian elephants (*Elephas maximus*), we assessed the relative virulence $v_{\mathrm{J}}/v_{\mathrm{A}}$ from the published literature (Lynsdale *et al.* 2017) as well as personal communication with C. Lynsdale and V. Lummaa. The censused individuals (in total 4242) are categorized into reproducible (aged 8 or above; 3046 individuals) – or adults – and nonreproducible (under 8; 1196 individuals) – or juveniles (Sukumar *et al.* 1997). Parasite-caused and potentially parasite-associated death, in total, occurred in 304 adults or in 301 juveniles, each among which parasite-caused death was identified as 85 for adults or 91 for juveniles, respectively (we thank C. Lynsdale and V. Lummaa for sharing the data of stage-specific mortality). From these data, however, we were unable to assess the virulence values per se (because we lack data for stage-specific prevalence or proportion of infected individuals among those at the same stage). Therefore, we restricted ourselves to citing the evidence that extremely young individuals are at higher risk of parasite-induced death (see Figure2 in Lynsdale *et al.* 2017). We propose that future studies quantifying stage-dependent parasite prevalence is greatly promising to test our predictions.

All drawings were downloaded from PhyloPic. Credits: (a) Uncredited; (b) David Liao, under CC BY-SA 3.0; (c, d) Both uncredited; (e) T. Michael Keesey, under CC BY 3.0 (the image has been reflected from original); (f) Uncredited; (g) Anthony Caravaggi, under CC BY-NC-SA 3.0 (the image has been reflected from original); (h) Luc Viatour (source photo) and Andreas Plank, under CC BY-SA 3.0.

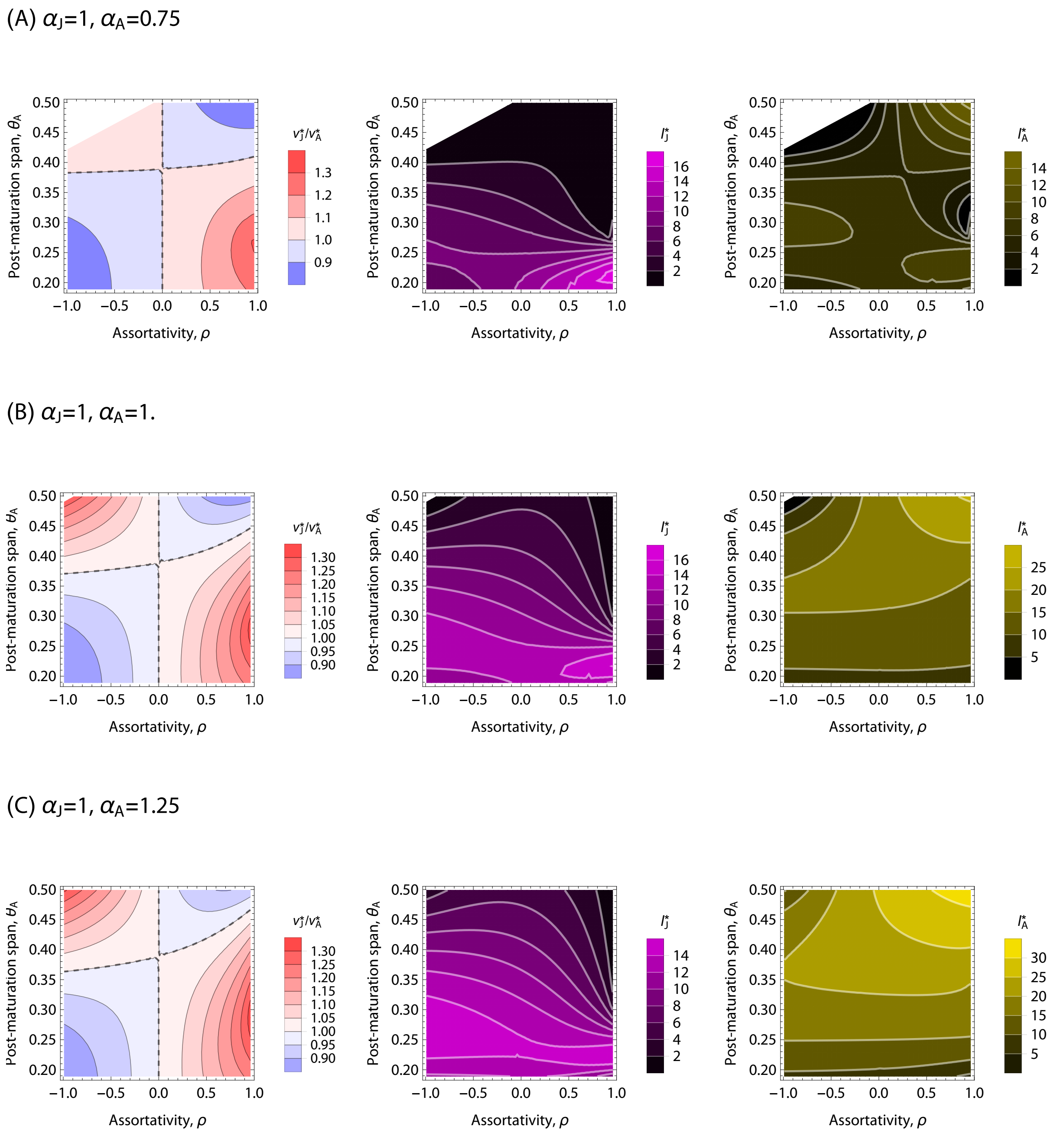

SI Figure 3: Effects of varying susceptibility. Changes in susceptibility have minor effects on the CSS (left panels), whereas evolutionary suicide is more likely to occurs with smaller suceptibility (panel A's).

(A) $\tau_J$=0.375, $\tau_A$=0.

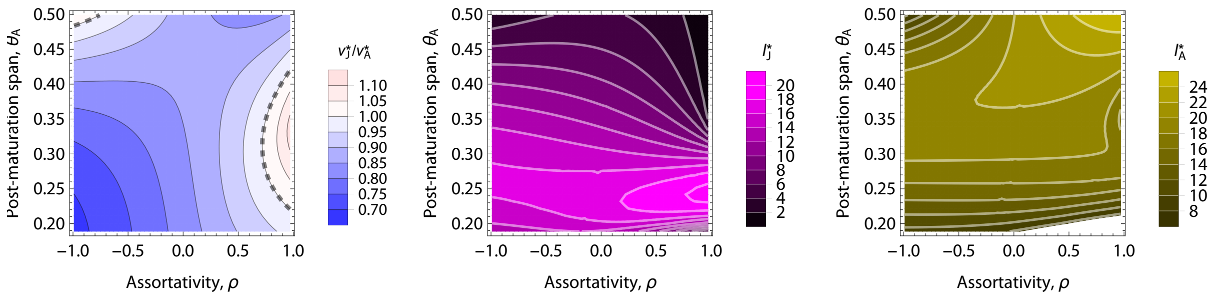

(B) $\tau_J$=0, $\tau_A$=0.375

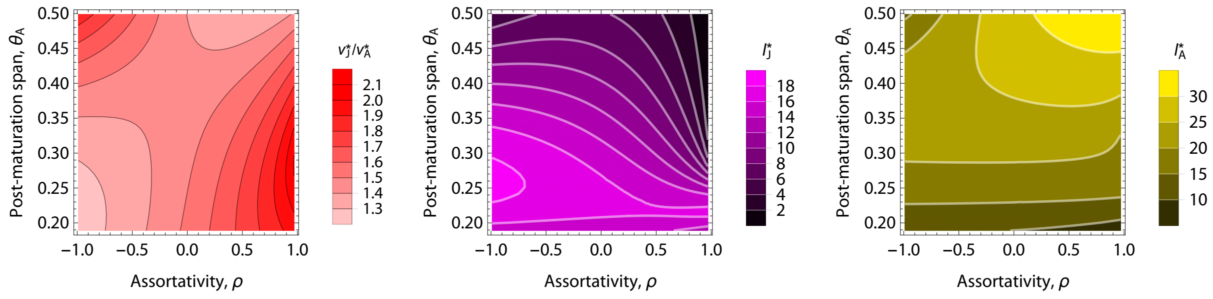

SI Figure 4: Effects of varying tolerance. Tolerance in adults can lead to relatively higher virulence for juveniles; note that $v^*_A = \sqrt{m_A(1-\tau_A)/k_A}$ is dependent on $\tau_A$. Due to the tolerance, the number of infected adults increase with $h_A$. Overall, the qualitative trend is unchanged.

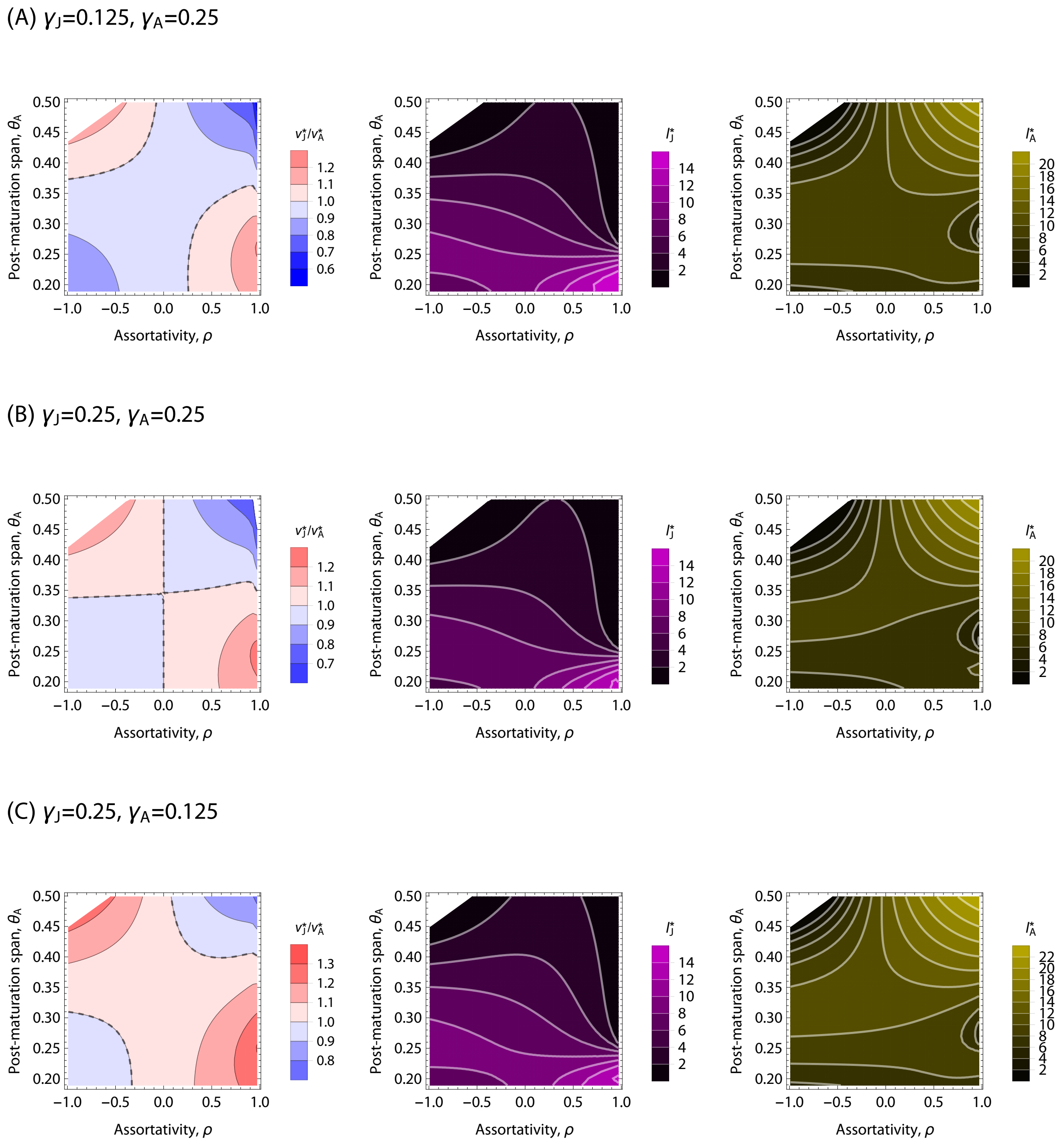

SI Figure 5: Effects of varying recovery rates. The results are quantitatively unchanged, but evolutionary suicide is more likely to occur (white zone). Dashed contours: $v_J^* = v_A^*$. Default values were used for other parameters (main text).

(A) $b_J$=10, $b_A$=8.

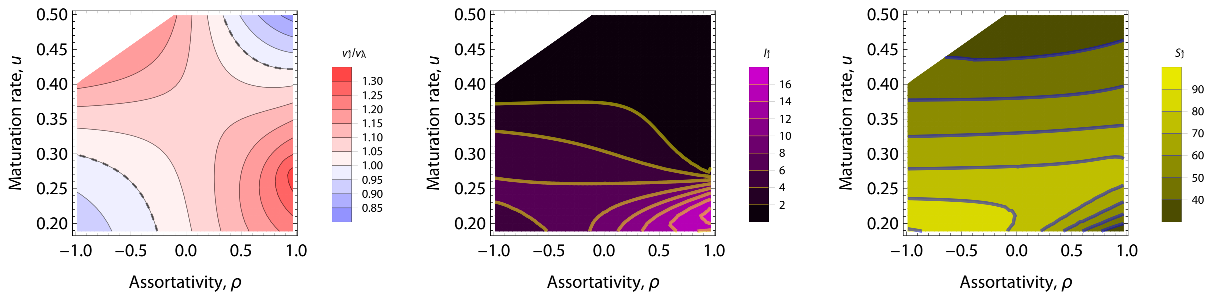

(B) $b_J$=10, $b_A$=14.

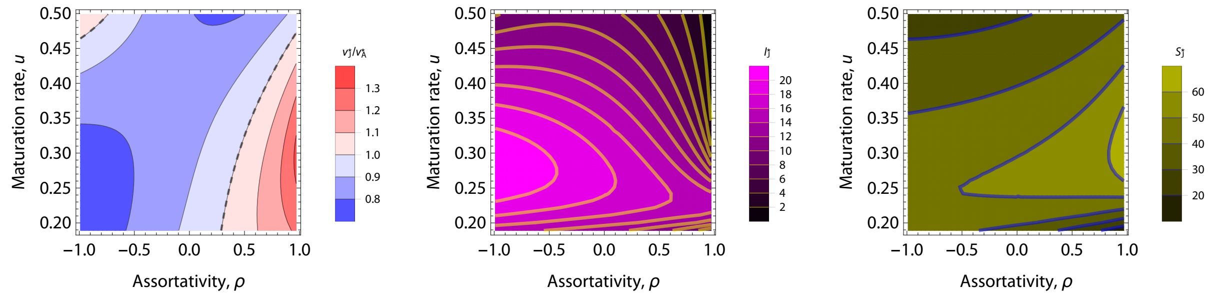

SI Figure 6: Effects of varying infectiousness. Different infectiousness can lead to higher virulence for juveniles; note that $v_A^* = \sqrt{m_A/k_A}$ is independent of $b_A$ and $b_J$. Overall, the qualitative trend is unchanged, but the disease prevalence among juveniles is dramatically lower with assortativity.

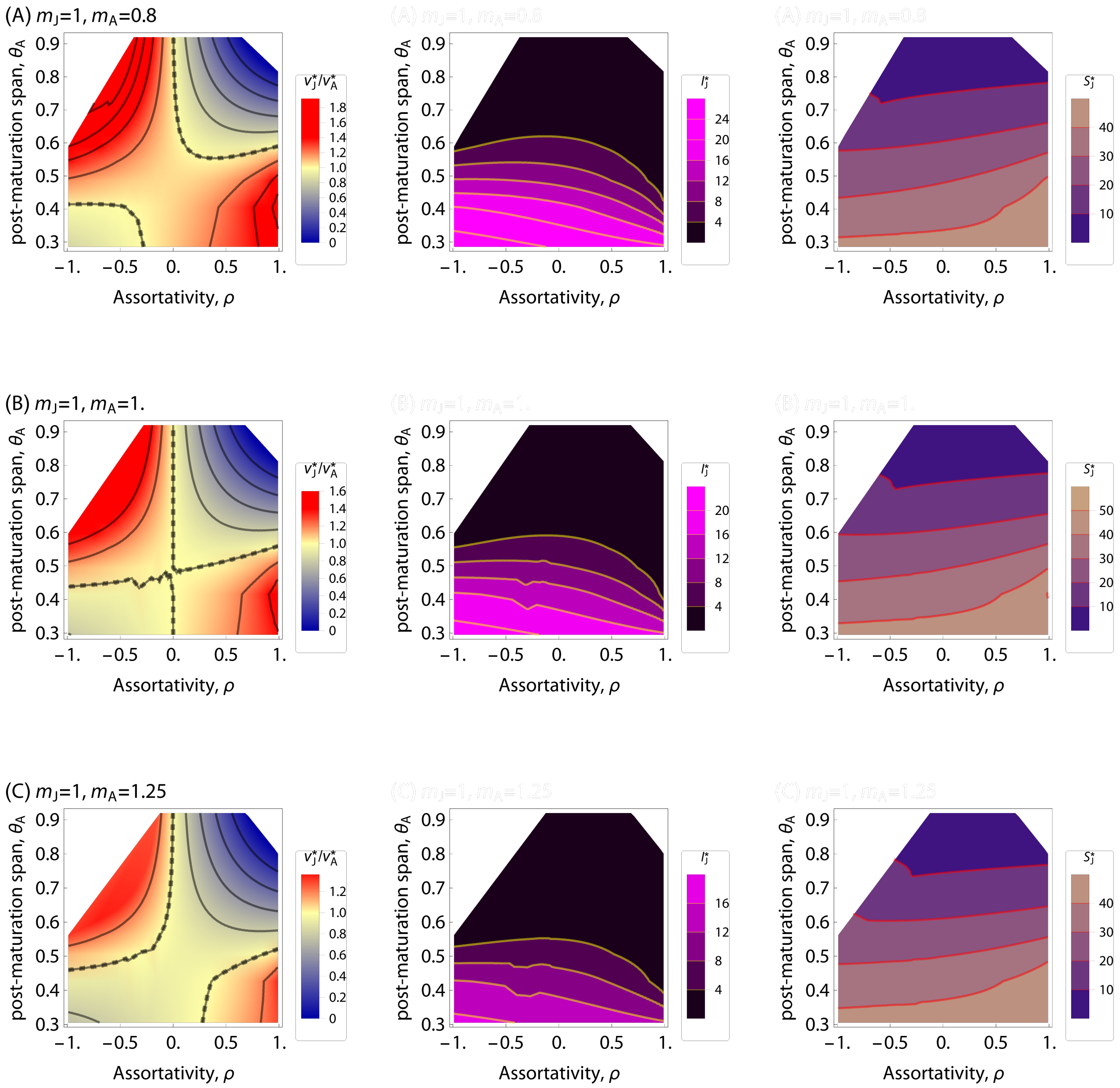

SI Figure 7: Effects of density-dependence.

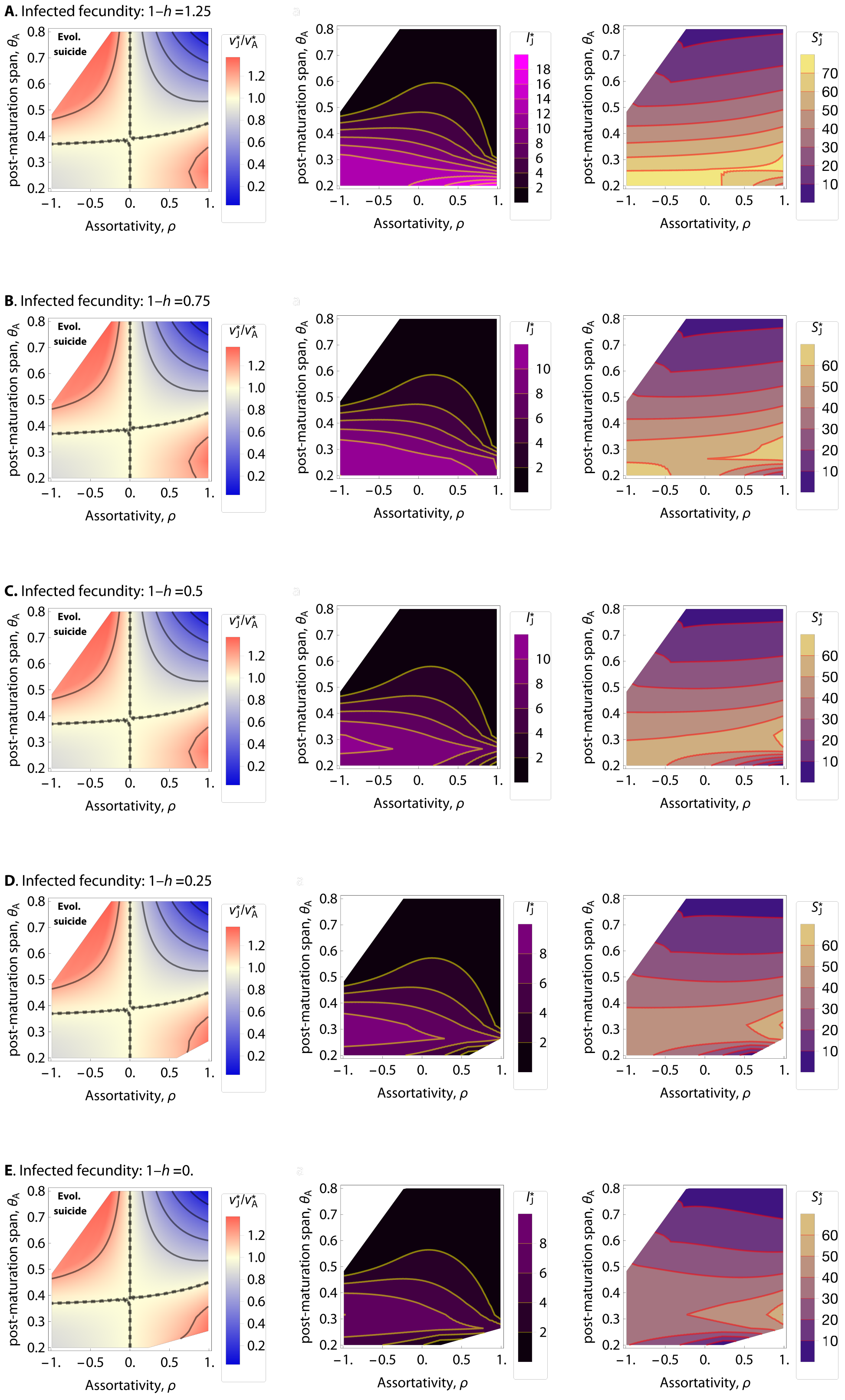

SI Figure 8: The effects of constant fecundity virulence, with $1-h$ measuring the fecundity of infected adults. The resulting difference is minor, as fecundity reduction acts only via ecological feedback without any direct effects on the invasion fitness. Also note that in panel (A), the fecundity is higher for infected than for susceptible adults.

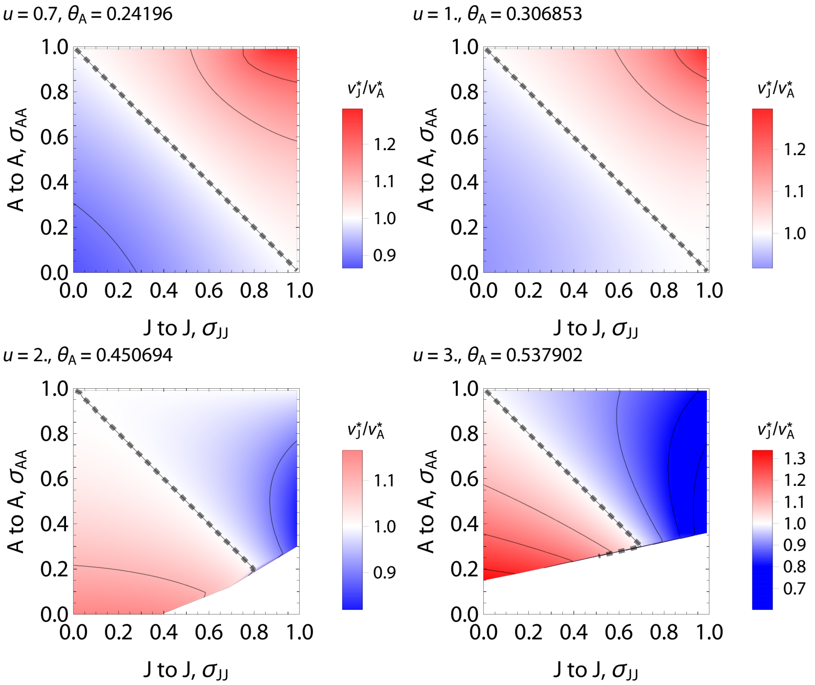

SI Figure 9: Normalized pathway structure, with $\sigma_{\mathrm{JA}} = 1 - \sigma_{\mathrm{AA}}$ and $\sigma_{\mathrm{AJ}} = 1 - \sigma_{\mathrm{JJ}}$. Orthogonal dashed line, which satisfies $\rho = \sigma_{\mathrm{JJ}} + \sigma_{\mathrm{AA}} - 1 = 0$, gives $v_{\mathrm{J}}^* = v_{\mathrm{A}}^*$. Note, we fixed $m_{\mathrm{J}} = m_{\mathrm{A}} = 1$, and thus $\theta_{\mathrm{A}}$ is a function of $u$ (e.g., $u = 1$ gives $\theta_{\mathrm{A}} = 0.306853$).

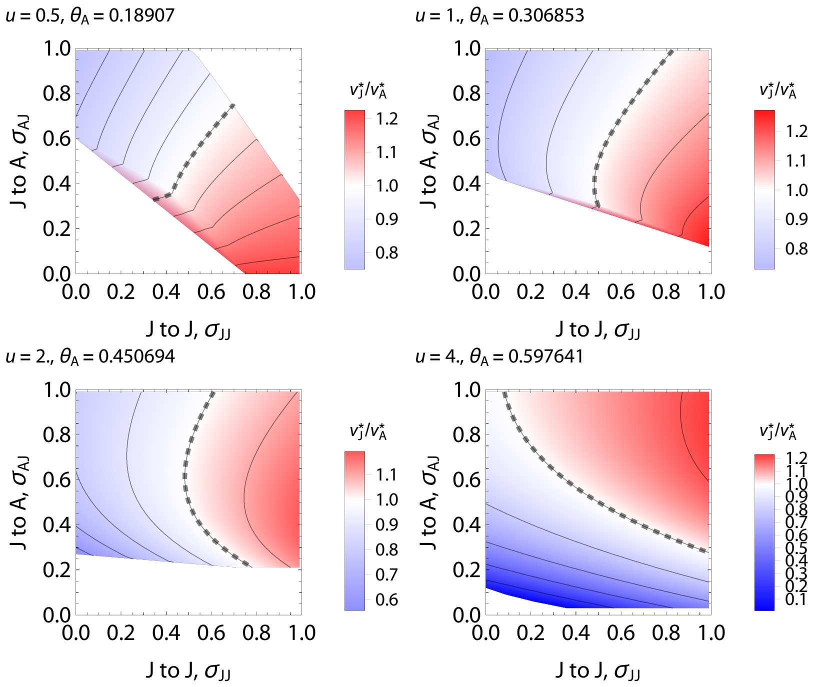

SI Figure 10: Symmetric pathway structure, with $\sigma_{\mathrm{JA}} = \sigma_{\mathrm{AJ}}$ varied and $\sigma_{\mathrm{AA}} = 0.5$ fixed. Note that fixing $u$ determines a single value of $\theta_{\mathrm{A}}$, and for clarity we have shown both of the values ($u$ and $\theta_{\mathrm{A}}$).

SI Table: Empirical data on empirical host-parasite systems.

| Host | Parasites | vJ/vA | Adult Period | Assorativity | Refs |
|---|---|---|---|---|---|
| **Pink salmon (*Oncorhynchus gorbuscha*)** | Salmon Louse | >1 | ≈0 (semelparity) | Assortative | Heard 1991; Jones *et al*. 2008 |
| **Gerbil (*Gerbillus andersoni*)** | Ectoparasites | 1.79 | 0.41 | – | Wassif & Soliman 1980; Delany 1986; Hawlena *et al.* 2006 |
| **Fruit fly (*Drosophila melanogaster*)** | Bacteria (*Pseudomonas entomophila*) | 1.43 | 0.63 | – | Vodovar *et al.* 2005; Luckinbill *et al*. 1984 |
| **Common guillemot (*Uria aalge*)** | Great Island Virus | 0.69 | 0.70 | Assortative | Harris & Wanless 1995; Nunn *et al*. 2006; Wanelik *et al*. 2017 |
| **Asian elephant (*Elephas maximus*)** | Parasites | >1 | 0.76 | – | Sukumar *et al*. 1997; Lynsdale *et al*. 2017 |
| **European rabbit (*Oryctolagus cuniculus*)** | Nematode | 1.00 | 0.83 | – | von Holst *et al*. 2002; Cornell *et al*. 2008 |
| **Rabbits (*Leporidae*)** | RHD Virus | 0.67 | 0.83 | – | Morisse *et al*. 1991; Reluga *et al*. 2007 |
| **Pigeon (*Columba livia*)** | Blood Parasite | 1.85 | 0.92 | – | Lack 1968; Holmes & Ottinger 2003; Sol *et al*. 2003 |
